# Hypoxia-driven deSUMOylation of EXOSC10 promotes adaptive changes in the transcriptome profile

**DOI:** 10.1101/2023.06.23.546038

**Authors:** Chrysa Filippopoulou, Chairini C. Thome, Sofia Perdikari, Evgenia Ntini, George Simos, Katherine E. Bohnsack, Georgia Chachami

## Abstract

Reduced oxygen availability (hypoxia) triggers adaptive cellular responses via hypoxia-inducible factor (HIF)-dependent transcriptional activation. Adaptation to hypoxia also involves transcription-independent processes like post-translational modifications, however these mechanisms are poorly characterized. Investigating the involvement of protein SUMOylation in response to hypoxia, we discovered that hypoxia strongly decreases the SUMOylation of Exosome subunit 10 (EXOSC10), the catalytic subunit of the RNA exosome, in a HIF-independent manner. EXOSC10 is a multifunctional exoribonuclease enriched in the nucleolus that mediates the processing and degradation of various RNA species. We demonstrate that the Ubiquitin-specific protease 36 (USP36) SUMOylates EXOSC10 and we reveal SUMO1/sentrin-specific peptidase 3 (SENP3) as the enzyme mediating deSUMOylation of EXOSC10. Under hypoxia, EXOSC10 dissociates from USP36 and translocates from the nucleolus to the nucleoplasm concomitant with its deSUMOylation. Loss of EXOSC10 SUMOylation does not detectably affect rRNA maturation but affects the mRNA transcriptome by modulating the expression levels of hypoxia-related genes. Our data suggest that dynamic modulation of EXOSC10 SUMOylation and localization under hypoxia regulates the RNA degradation machinery to facilitate cellular adaptation to low oxygen conditions.

## 1. Introduction

Oxygen is indispensable for the survival of multicellular organisms as it is essential for mitochondrial bioenergetics and numerous biochemical reactions. When oxygen requirements exceed its supply (a condition known as hypoxia), cells rapidly adapt their gene expression programs to reduce oxygen consumption and increase anaerobic energy production (1,2). In cancer, hypoxia is a hallmark of the tumor microenvironment that affects the metastatic potential of tumor cells and the behavior of stromal cells (3).

A key aspect of the adaptive response to hypoxia at the transcriptional level is activation of a family of heterodimeric transcription factors called hypoxia inducible factors (HIFs) (4,5). HIF transcriptional activity depends on the oxygen-dependent stabilization of the HIF-α subunit. Typically, HIF-regulated genes encode proteins involved in processes such as angiogenesis, erythropoiesis, autophagy, lipid and glucose metabolism, invasion and metastasis (6,7). Notably however, adaptation to hypoxia also involves transcription-independent processes that are less characterized. These include rearrangements of the actin cytoskeleton (8), changes in mRNA translation (9) and alternative splicing (10). It was also shown that short-term exposure to hypoxia causes nuclear matrix and splicing machinery restructuring via a reactive oxygen species (ROS)-dependent mechanism (11). Hypoxia can, under certain circumstances (e.g. when combined with acidosis), also affect nucleolar architecture and ribosome biogenesis (12). Interestingly, the nucleolus acts as a stress sensor and signaling hub under various harmful conditions such as nutrient deprivation, DNA damage and oxidative or heat stress (13,14). Many of these post-transcriptional responses are mediated by post-translational protein modifications.

SUMOylation, the covalent attachment of SUMO proteins to target proteins, has been recently shown to play a role in the response to hypoxia as both HIFs and HIF-targets can be directly modified with SUMO (15). SUMOylation is a dynamic process and contributes to the regulation of transcription, protein localization and other processes under stress conditions (16–19). Similar to the ubiquitin system, SUMO conjugation to target proteins requires specific E1, E2 and E3 enzymes. The E2 SUMO conjugating enzyme, namely UBC9, can transfer SUMO to its target protein either with or without the involvement of an E3 ligase (20). UBC9 typically recognizes a ψKxD/E motif (ψ: aliphatic branched amino acid, K: target lysine, x: any amino acid, D/E: aspartate or glutamate) within the substrate protein, although SUMOylation can also occur on lysine residues not within the consensus motif (21). Notably, SUMO recruitment to target lysine residues can be facilitated by nearby SUMO-interacting motifs (SIM). A SUMO E3 ligase is normally required to cooperate with UBC9 for efficient covalent attachment of SUMO to target proteins. So far, only few proteins have been characterized as SUMO E3 ligases through their ability to enhance SUMOylation of targets, and these include proteins of the protein inhibitor of activated STAT (PIAS) family, Topors, RanBP2, Pc2, p14^ARF^, Krox20, SF2/ASF and ZNF451(22–27). The nucleolar protein USP36, which is a well-characterized deubiquitinating enzyme (DUB), was recently suggested to be an E3 SUMO ligase, promoting the SUMOylation of small nucleolar ribonucleoprotein (snoRNP) components (28). SUMOylation can be reversed by the action of SUMO-specific isopeptidases, the biggest family of which is the SUMO/sentrin-specific proteases (SENP) comprising SENP1, 2, 3, 5, 6, and 7 (29,30).

We previously combined SUMO-immunoprecipitation (IP) with quantitative mass spectrometry to identify SUMOylated proteins the SUMOylation status of which was altered by hypoxia without detectable concomitant changes in their protein expression levels (31). In addition to transcription factors, such as TFAP2A that could enhance the transcriptional activity of HIF-1 (31), the identified inventory of proteins included the Exosome component 10 (EXOSC10) (31), a well-characterized 3′-to-5′ exoribonuclease and part of the RNA exosome (32). The RNA exosome is the key RNA processing and decay machinery in human cells. It consists of nine core subunits (EXOSC1-9) arranged as a capped barrel-shaped catalytically inactive core, associated with two active subunits, the ribonucleases DIS3/DIS3L1 and EXOSC10 (33–36). EXOSC10, which localizes predominantly inside the nucleus and is enriched in the nucleolus, is involved, as part of the RNA exosome, in the processing, surveillance/quality control and degradation of various different RNA types (37). For example, EXOSC10 is implicated in 3’ end processing of ribosomal (r)RNA and small nuclear/nucleolar (sn/sno)RNA precursors (38–42). EXOSC10-associated RNA exosomes are also involved in the turnover of excised pre-rRNA spacer fragments and various non-coding RNA species, including long-non-coding (lnc)RNAs and cryptic unstable transcripts (CUTs) as well as the degradation of aberrant pre-RNAs, transfer (t)RNAs, messenger (m)RNAs, etc.(40,42–45).

Although our previous work as well as high-throughput studies have identified EXOSC10 as a target of SUMOylation (33,48–50), the role of this modification in the EXOSC10 function or the cellular response to hypoxia have not been previously addressed. Here, we report the identification of the enzymes responsible for EXOSC10 SUMOylation, and deSUMOylation, and describe how these processes and EXOSC10 function respond to low oxygen conditions. Together our data reveal the involvement of post-translational regulation of the RNA degradation machinery in the cellular adaptation to hypoxia.

## 2. Materials and Methods

### Plasmids

Human EXOSC10 cDNA was purchased from Gateway Full ORF clones of DKFZ, Heidelberg, Germany. The EXOSC10 coding sequence (NM_001001998.3) was cloned into the pcDNA 3.1-HA vector (kindly provided by Melchior F., ZMBH, University of Heidelberg, Germany) as BamHI-XbaI fragment. For the expression of siRNA-insensitive (siins) EXOSC10, the EXOSC10 cDNA containing silent mutations in the siRNA target sites was purchased in a pEX-A258 vector from Eurofins (Germany) and cloned into the pcDNA5-2xFlag-His_6_ (Flag) vector as a BamHI-NotI fragment. Site-directed mutagenesis was used to introduce point mutations in the wild-type (WT) sequences of pcDNA 3.1-HA-EXOSC10 and pcDNA5-2xFlag-His_6_-siinsEXOSC10 to generate mutants encoding EXOSC10 versions with specific lysine to arginine substitutions (K583R, K168R, K201R). For bacterial overexpression of GST-EXOSC10 WT, K583R and D313A fusion proteins, the EXOSC10 coding sequence was cloned from pcDNA 3.1-HA-EXOSC10 plasmids to pBluescript II SK (+/-) as a BamHI-XbaI fragment and then subcloned as a BamHI-NotI fragment in pGEX-4T1 bacterial expression vector. See Supplementary Table S1 for cloning and site-directed mutagenesis primers and Supplementary Table S2 for a list of plasmids used in this study.

### Cell culture and cell treatments

Human HeLa cells, HEK293 Flp-In T-REx cells (Thermo Fischer Scientific, Waltham, MA, USA) and HeLa Flp-In T-REx cells (kindly provided by Mayer T., Universität Konstanz, Germany) were cultured at 37°C with 5% CO_2_ in Dulbecco’s modified Eagle’s medium supplemented with 10% fetal bovine serum and penicillin–streptomycin (Biosera, Nuaille, France). For hypoxic treatment, cells were exposed to 1% O2, and 5% CO2 in an INVIVO2 200 hypoxia workstation (Ruskinn Life Sciences, Pencoed, UK) for the indicated times. When required, cells were treated with 2 μm rapamycin for 6 h (Sigma-Aldrich, St Louis, MO, USA) or DMSO at the appropriate concentration as a solvent control.

HeLa Flp-In T-REx cells were transfected with pcDNA5-2xFlag-His6-siinsEXOSC10 (WT or K583R) or pcDNA5-2xFlag-His6 (empty vector) plasmids using XtremeGene9® transfection reagent (Sigma Aldrich, St Louis, MO, USA) according to the manufacturer’s guidelines. Transfected cells in which the transgene was integrated into the Flp-In locus were selected with 1 µg/mL of blastocidin and 200 µg/mL of hygromycin.

### Cell transfections for siRNA-mediated gene silencing and protein expression from plasmid DNAs

For gene silencing, cells were transfected with the appropriate siRNAs (see Supplementary Table S3) using RNAiMax reagent (Invitrogen, Life Technologies, Carlsbad, CA, USA) according to the manufacturer’s protocol and harvested after 48 h or 72 h. AllStars siRNA (Qiagen, Venlo, Netherlands) or an siRNA targeting firefly luciferase (46) (see Supplementary Table S3) were used as negative control. For experiments using the rescue system, the cells were transfected with appropriate siRNAs (see Supplementary Table S3) and harvested after 72 h. 24 h prior harvesting, cells were treated with 1 µg/mL tetracycline/doxycycline to induce expression of 2xFlag-His6-siinsEXOSC10 (WT or K583R) or the 2xFlag-His6-tag only. Cells were transfected with plasmid DNAs using polyethylenimine (PEI, Thermo Scientific, Waltham, MA, USA).

### Protein extraction, SDS-PAGE, western blotting and antibodies

Proteins were extracted from cells using radioimmunoprecipitation assay (RIPA) buffer followed by protein precipitation with trichloroacetic acid (TCA). Proteins were resolved on denaturing (sodium dodecyl sulphate (SDS)) 7%–15% or gradient 4-20% polyacrylamide gels and subjected to SD-PAGE electrophoresis, transferred to nitrocellulose membrane and analyzed by western blotting using antibodies listed in Supplementary Table S4. Western blot images were taken by using a Uvitec Cambridge Chemiluminescence Imaging System supplied with ALLIANCE SOFTWARE (ver. 16.06) and quantified by UVIBAND SOFTWARE (ver. 15.03) supplied by the instrument manufacturer (Uvitec Cambridge, Cambridge, UK).

### Endogenous SUMO-1 immunoprecipitation (IP) and non-denaturing-IP

Endogenous SUMO-1 IP from HeLa or HEK293 cells was performed as described previously (47,48). Briefly, cells were lysed using a denaturing lysis buffer with 1% SDS, the lysates were diluted 10-fold to reach a final SDS concentration of 0.1% and were incubated with monoclonal anti-SUMO1 (SUMO1 21C7)-coupled beads at 4 °C overnight. For non-denaturing IP conditions, cells were lysed in buffer containing 20 mM Tris pH 7.4, 150 mM NaCl, 0.5% Triton X-100, 1mM EDTA, 50 mM glycerolphosphate, 10 mM Na_3_VO_4_, 20 mM N-Ethylmaleimide, supplemented with Protease Inhibitor Mix G (SERVA Electrophoresis GmbH, Germany). Cell lysates were incubated for 3 h with anti-EXOSC10 and for 1 additional hour in the presence of Protein G-Sepharose beads. Bound proteins were eluted in SDS sample buffer [50 mM Tris pH 6.8, 2% SDS, 0.1% bromophenol blue, 10% glycerol, 50 mM Dithiothreitol (DTT)]. For non-denaturing IP conditions from nuclear extracts, HeLa cells were resuspended in Cytoplasmic Lysis Buffer containing 10 mM Tris pH 8, 10 mM NaCl, 1.5 mM MgCl_2_, 20 mM N-Ethylmaleimide, supplemented with Protease Inhibitor Mix G for 30 min, NP-40 was added to final concentration of 0.3% and cells were homogenized using a precooled glass Dounce Homogenizer. Purified nuclei were pelleted by centrifugation at 1200 x g for 5 min, washed once with Cytoplasmic Lysis Buffer, resuspended in Nuclear IP buffer containing 50 mM Tris pH 8, 150 mM NaCl, 0.5% NP-40, 1mM EDTA, 50 mM glycerolphosphate, 10 mM Na_3_VO_4_, 20 mM N-ethylmaleimide, supplemented with Protease Inhibitor Mix G and sonicated for 25 sec (0.5 sec pulses). Nuclear extracts were cleared by centrifugation at 15,000 x g for 15 min and incubated for 12 h with anti-EXOSC10 and for 1 additional hour in the presence of Protein G-Sepharose beads. Bound proteins were eluted in SDS sample buffer.

### Immunofluorescence and Image analysis

HeLa cells were grown on coverslips, incubated in normoxia or hypoxia for 24 h and prepared for immunofluoresence microscopy as previously described (49,50) using the indicated primary antibodies (see Supplementary Table S4), and Alexa 488-or 594-conjugated anti-rabbit or anti-mouse secondary antibodies (1: 1000, Jackson ImmunoResearch, Cambridgeshire, UK). For studying the localization of 2xFlag-His_6_-EXOSC10 WT and K583R proteins, HeLa Flp-In cells were grown on coverslips, treated with 1 μg/ml doxycycline for 30 h (6 h prior to incubation in normoxic or hypoxic conditions) and processed as described above using an anti-Flag primary antibody. Cells were visualized in a Zeiss Confocal microscope (Zeiss, Oberkochen, Germany) or a Zeiss Axio Imager.Z2 microscope. Scale bars were set to 10μm or 1μm.

All images were processed with Zen software (2011, blue edition, Zeiss Oberkochen, Germany) and FiJi-ImageJ (NIH, Bethesda, MD, USA). For representation, images were saved as 32-bit color images. For quantification of nucleolar and nucleoplasmic signal, cells were captured with the same exposure time and the original unmodified images (8-bit) were used with no other adjustment. After subtracting background by setting rolling ball diameter on 50-pixel, cell areas (total, nuclear, nucleolar) were manually selected and thresholded, then area, mean gray value of background and integrated density were calculated for the selected regions of interest. Corrected total cell fluorescence (CTCF) is presented in the quantitative graphs as calculated by CTCF= integrated density – (selected area × mean gray value of background).

### Protein purification and in vitro sumoylation assays

Plasmids encoding glutathione-S-transferase (GST) -tagged EXOSC10 WT/K583R/D313A or GST-USP36 1-420/421-800 were used for transformation of the BL21(RIL) strain of *E. coli*. Overexpression and purification of the GST-tagged EXOSC10 forms and GST-tagged USP36 forms was performed as previously described (28,38). Briefly, protein expression of GST-tagged EXOSC10 was induced by adding 1 mM isopropyl β-D-1-thiogalactopyranoside (IPTG) for 16 h at 4 °C. Bacterial cells were then, lysed by sonication in lysis buffer containing 50 mM Tris-HCl pH 8.0, 10 % glycerol, 300 mM NaCl, 5 mM MgCl_2_, 0.1 % Tween20 supplemented with Protease Inhibitor Mix G (SERVA Electrophoresis GmbH, Germany). For the GST-tagged USP36 versions, protein expression was induced by adding 0.5 mM IPTG for 4 h at 25 °C and bacterial cells were lysed by sonication in lysis buffer containing 20 mM HEPES pH 7.5, 150 mM NaCl, 5 mM MgCl_2_, 1% Triton X-100, 1 mM DTT supplemented with Protease Inhibitor Mix G (SERVA). GST-tagged proteins in soluble extracts were purified by GSH-agarose beads (Macherey-Nagel, Düren, Germany) from which were eluted by the addition of 50 mM reduced L-glutathione (dissolved in 25 mM Tris-HCl pH 8).

*In vitro* sumoylation assays were previously described (51). Reactions containing 0.25 μM of GST-tagged EXOSC10 WT/K583R and all the necessary purified SUMOylation machinery components (kindly provided by F. Melchior lab, University of Heidelberg, Germany): untagged hSUMO1 (9 μΜ), untagged E2 (Ubc9) (as indicated), E1 (His-Aos1/untagged Uba2) (70nM) and either His-RanBP2-BD3-4 (aa 2304-3062) (100nM), GST-USP36 1-420 (100nM) or GST-USP36 421-800 (100nM), diluted in SUMOylation assay buffer (20 mM HEPES pH 7.3, 110 mM potassium acetate, 2 mM magnesium acetate, 1 mM EGTA, 1 mM DTT, 0.05% Tween 20, 0.2 mg/ml ovalbumin, Protease Inhibitor Mix G) were performed for 2 h or 4 h at 30 °C in the presence or the absence (control reactions) of ATP.

### In vitro 3’-5’ exoribonuclease assay

The in vitro exoribonuclease assay was performed as previously described (38) using 400 fmol of recombinant GST-EXOSC10 (WT, K583R or D313A) and 10 fmol of 5’-[32P]-labeled poly-uridine (U32) RNA substrate. The recombinant proteins were pre-incubated at 37°C for 10 min in a buffer containing 10 mM Tris–HCl pH 7.6, 75 mM NaCl, 2 mM DTT, 100 µg/ml bovine serum albumin (BSA), 0.8 U/µl RiboLock (Thermo Fisher Scientific), 4.5 % glycerol, 0.05 % Triton X-100, 0.5 mM MgCl2, after which, the 5’-[32P]-U32 was added and samples were further incubated at 37°C. Aliquots were collected after 0, 10, 30, 60 and 120 minutes and the reaction was stopped by addition of 2x RNA loading-dye (80 % formamide, 10 mM EDTA pH 8.0, 1 mg/ml xylene cyanol FF, 1 mg/ml bromophenol blue). RNA samples were separated on a denaturing (7 M urea) 15% polyacrylamide gel in 0.5X Tris-borate-EDTA (TBE) buffer at 20 W. The gel was dried and exposed to a phosphorimager screen. Images were later obtained using a Typhoon FLA 9500 (GE).

### RNA extraction, RNA electrophoresis and northern blotting

Total RNA was extracted using TRI reagent (Sigma-Aldrich) according to the manufacturer’s instructions. For analysis of high-molecular-weight RNA species, 4 µg of total RNA were denatured in 5 volumes of Glyoxal-loading dye (61% dimethyl sulfoxide (DMSO), 20% glyoxal, 1X BPTE [30 mM Bis-Tris, 10 mM piperazine-N,N′-bis(2-ethanesulfonic acid) (PIPES), 1 mM EDTA], 5% glycerol) for 1 h at 55°C and separated in a 1.2% agarose-BPTE gel for 16 h at 60 V. After washing the gel with 100 mM NaOH for 20 min, twice in a buffer containing 0.5 M Tris pH 7.4 and 1.5 M NaCl for 15 min, and once in 6X saline sodium citrate (SSC; 0.9 M sodium chloride, 90 mM sodium citrate) buffer for 15 min, the RNAs were transferred to Hybond-N+ membrane by vacuum blotting. For low-molecular weight RNA species, 2.5 µg of total RNA were separated in a denaturing (7 M urea) 10% polyacrylamide gel in 0.5X TBE buffer and transferred to Hybond-N+ membrane by electroblotting.

After the RNAs were crosslinked to membranes with UV-light (254 nM), the membranes were pre-incubated with hybridization buffer (250 mM sodium phosphate pH 7.4, 7% SDS and 1 mM EDTA pH 8.0) for 30 min and hybridized with [32P]-labeled DNA oligonucleotide probes (see Supplementary Table S1). Membranes were then washed with 6X SSC and 2X SCC supplemented with 0.1 % SDS each for 30 min at 37°C and exposed to phosphorimager screens. Images were obtained with Typhoon FLA 9500 (GE) and quantified using the Image Studio 5.2.5 software (LI-COR).

### Real-time quantitative PCR (qPCR)

500 ng of total RNA extracted using TRI reagent (Sigma-Aldrich) was reverse transcribed using PrimeScript™ RT Reagent Kit (Perfect Real Time) (Takara Bio, Kusatsu, Japan) according to the manufacturer’s instructions. 10μl of cDNA synthesis reactions were performed, containing 1x PrimeScript Buffer, PrimeScript RT Enzyme Mix I and anchored oligo d(T) primer in the following conditions: 5 min at 25 °C, 15 min at 37 °C, 5 sec at 85 °C. cDNAs were diluted in 1:10 ratio and used for qPCR analyses. Specific forward and reverse primer pairs (intron spanning) for the amplification of the target mRNAs were designed to generate amplicons of 70-100 bp and have equal melting temperatures (See Supplementary Table S5). The efficiency of the primers was validated by checking for amplification of a single amplicon using a product melting curve and the resulting PCR-products were visualized in 2% agarose gels containing ethidium bromide (EtBr). The amplification efficiency and linearity of cDNA amplification was determined and only primer pairs with an amplification efficiency of >90% were used. qPCR was performed using KAPA SYBR FAST qPCR (Kapa Biosystems, Wilmington, MA, USA) and 200 nM primers in a LightCycler 96 (Roche Life Science, Basel, Switzerland). Cycling conditions were 95 °C for 3 min, 40 cycles of 95 °C for 10 sec, 60 °C for 20 sec and 72 °C for 1 sec, followed by 95 °C for 5 sec, 65 °C for 1 min and 97 °C for 30 sec. qPCR data were analyzed using LightCycler 96 system (version 1.1.0.1320). Two biological independent experiments for the rescue system and three biological independent experiments for the normoxia/hypoxia conditions were performed, and each sample was assayed in technical duplicates. Values differing by > 0.5 Cq were excluded. Relative quantitative gene expression was calculated using the DDCT method, normalized to RPLP1 mRNA levels and is presented as a fold increase in relation to the appropriate control condition (nt for the rescue system and normoxia for the normoxia/hypoxia conditions) according to the following formula:

2^Ct(control average of gene of interest)-Ct(gene of interest of each sample)^/2^Ct(control average of RPLP1)-Ct(RLPL1 of each sample)^

### RNA sequencing (RNA-seq)

Total RNA (10 µg) extracted using TRI reagent (Sigma-Aldrich) as described above was further purified using the RNA Clean&Concentrator (Zymo Research) kit according to the manufacturer’s instructions. Samples were then depleted of rRNA and converted to cDNA libraries using the TrueSeq® Stranded Total RNA kit (Illumina). cDNA libraries were sequenced with the HiSeq 4000 Sequencing System (Illumina) at the NGS Integrative Genomics Core Unit (NIG) at the University Medical Centre Göttingen, Germany. The quality of 50 basepair (bp) single-end reads was assessed using the FASTQC (v0.11.9) tool. Reads were aligned to the GRCh38.p13 human genome assembly (hg38) with annotation from Gencode (version 41) using STAR (v2.7.10a). Alignment allowed only 1 mismatch per read and reads were counted using --quantMode GeneCounts. DESeq2 (v1.36.0) was used to normalize the read counts and to calculate the differential expression of genes between two conditions. Principal Component Analysis was performed with *regularized log* (rlog) transformed count data. Rlog transformation of count data from all genes was done using the *rlog* function from DESeq2 package. Gene ontology analysis using the R package “clusterProfiler” (52) was performed on the DE gene lists generated by group comparisons to identify the enriched biological pathways. Ensembl based annotation for the target-genes and Genome wide annotation for Human ‘‘org.Hs.eg.db’’ were used. The raw RNA-seq data are deposited in Gene Expression Omnibus (GEO) database [http://www.ncbi.nlm.nih.gov/geo/] under the accession code GSE229894.

### Statistical analysis

Statistical differences were assessed using the GRAPH PAD PRISM version 6 software (GraphPad, San Diego, CA, USA). Data are expressed as mean SEM or mean SD as described. Differences were examined by ANOVA (one-way analysis–Tukey’s multiple comparisons) and Student’s t-test (two-tailed) where applicable. P < 0.05 was considered statistically significant (*P < 0.05; **P < 0.01; ***P < 0.001; ****P < 0.0001; n.s.: not significant).

## 3. Results

### 3.1 Hypoxia induces deSUMOylation of EXOSC10 in a HIF-independent manner

By combining SUMO-IP with quantitative mass spectrometry, we previously identified EXOSC10 as a SUMO-1 specific target, the SUMOylation of which was heavily reduced under prolonged hypoxia (48 h in 1% of O_2_) in HeLa cells (31). To confirm these findings in a non-cancer cell line and for a shorter exposure to hypoxia, the SUMO-1-IP was repeated using extracts from HEK293 cells incubated under normoxia or hypoxia for 24 h (Fig. 1A). SUMOylated proteins were precipitated in equal amounts from normoxic or hypoxic cells (Fig. 1B, bottom panel). SUMOylated EXOSC10 was readily detected, migrating approximately 20 kDa higher than the unmodified EXOSC10, in the SUMO-1 IP eluate from normoxic cells but was absent from the corresponding eluate from cells grown for 24 h under hypoxia, despite similar EXOSC10 protein expression levels (Fig. 1B, top panel, SUMOylated RanGAP1 is shown with an arrowhead and was used as a marker for equal loading in INPUTS). This indicates either inhibition of EXOSC10 SUMOylation or stimulation of EXOSC10 deSUMOylation in cells exposed to hypoxia.

**Figure 1:**
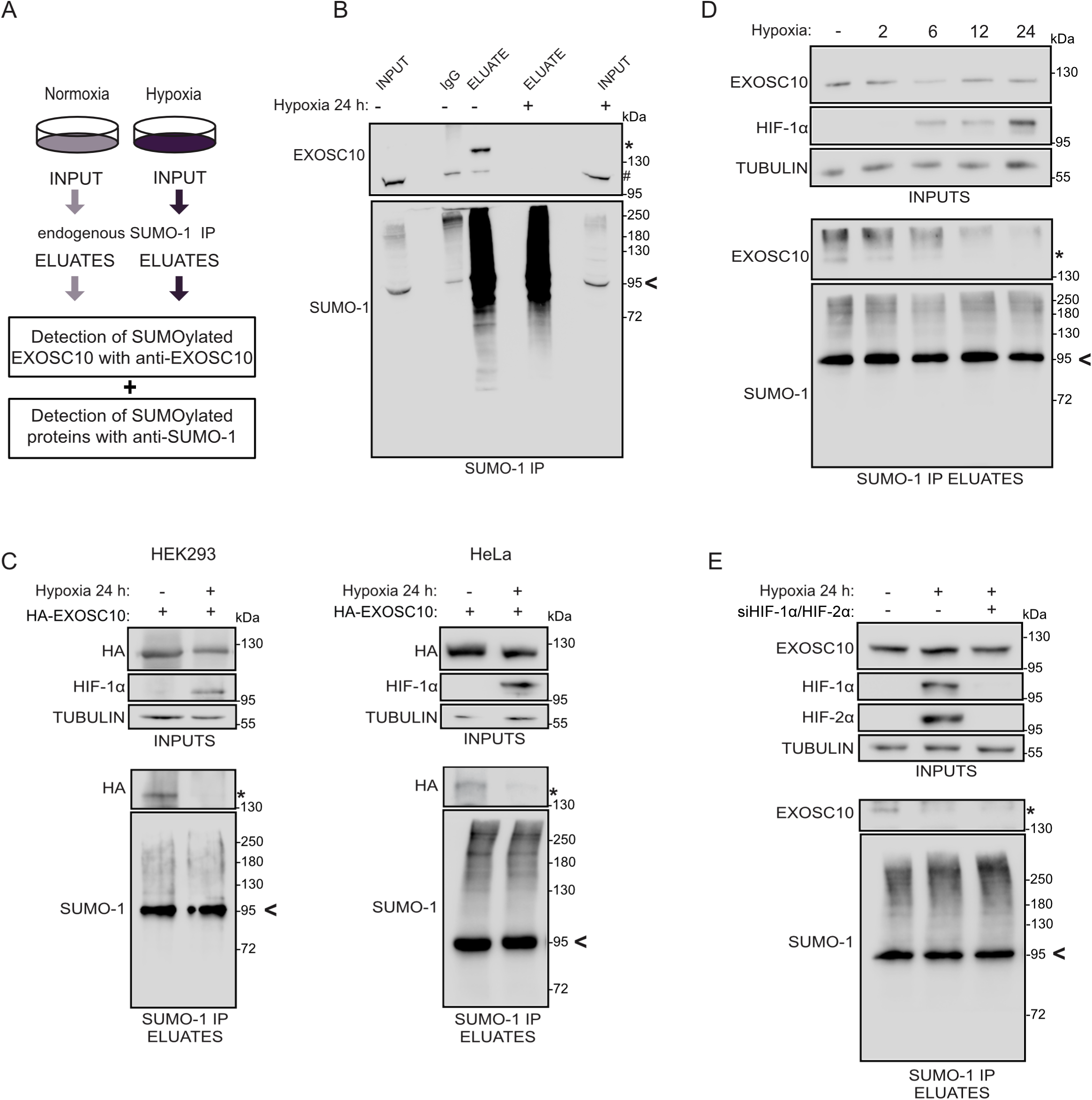
Hypoxia stimulates deSUMOylation of EXOSC10 in a HIF-independent manner. **A.** Schematic view of the anti-SUMO-IP used for the detection of EXOSC10 and other SUMOylated proteins. HEK293 cells were incubated in normoxia (-) or hypoxia (+, 1% of O_2_) for 24 h and their cleared lysates were subjected to SUMO-1 IP. **B.** Soluble extracts (INPUT), SUMO-1 and IgG immunoprecipitates (ELUATE), were analyzed by immunoblotting using the indicated antibodies. A rabbit anti-SUMO1 antibody was used for confirmation of endogenous SUMO-1 species enrichment. An equal amount of lysate was also loaded on IgG-beads and served as negative IP control. **C.** HEK293 (left) or HeLa cells (right) expressing the HA-EXOSC10 wild type construct were incubated in normoxia (-) or hypoxia (+) and their lysates were subjected to SUMO-1 IP and analyzed as described in (A). **D.** HeLa cells were incubated in normoxia (-) or hypoxia for the indicated times and their lysates were subjected to SUMO-1 IP and analyzed as described in (A). **E.** HeLa cells were transfected with a non-targeted siRNA (-) or siRNAs against HIF-1α and HIF-2α (+) and 24h post-transfection, were incubated in normoxia (-) or hypoxia (+) for 24h. Cell lysates were subjected to SUMO-1 IP and analyzed as described in (A). In all panels (A-D), the SUMOylated version of EXOSC10 is indicated with an asterisk (*). SUMOylated RanGAP1 is shown with an arrowhead and was used as a marker for equal loading in INPUTS (A) or as a marker for equal precipitation by anti-SUMO1. Beta-tubulin was used as loading control.

Further verification of these results was obtained by repeating the experiment with extracts from HEK293 and HeLa cells over-expressing an HA-tagged version of EXOSC10 incubated under either normoxia or hypoxia. Hypoxic conditions were verified by monitoring the levels of HIF-1α, which were markedly increased (Fig. 1C). SUMOylation of ectopically expressed EXOSC10 was almost abolished in both cell types specifically under hypoxia (Fig. 1C). Again, no strong differences in the protein expression levels of EXOSC10 under hypoxia were observed and this was also the case for other RNA exosome components, including the other catalytic subunit DIS3, the core proteins EXOSC2 and EXOSC3 or the associated factors C1D, MTR4 and MPP6, the expression levels of which were analysed in HeLa cells (Suppl. Fig. 1).

To test whether the reduction of EXOSC10 SUMOylation under hypoxia is linked to the key feature of response to hypoxia, namely the induction of HIFs and HIF-dependent transcriptional reprogramming, HeLa cells were exposed to hypoxia for different time points, between 2 and 24 h. Reduction of EXOSC10 SUMOylation was evident already after 2 h of hypoxic incubation while HIF-1α expression could only be detected after 6 h (Fig. 1D), suggesting that de-SUMOylation of EXOSC10 is an early event independent of HIF-regulated transcription. Consistent with this, reduced EXOSC10 SUMOylation was readily detected in hypoxic HeLa cells in which expression of HIF-1α and HIF-2α were effectively silenced with specific siRNAs (Fig. 1E).

### 3.2. Hypoxia induces nuclear redistribution of EXOSC10

To examine the effect of the hypoxia-induced de-SUMOylation on the biological function of EXOSC10, we first analysed its intracellular localization in HeLa cells. In normoxic cells, EXOSC10 was almost exclusively nuclear with prominent enrichment in the nucleoli as attested by its co-localization with two nucleolar markers, UBF (Upstream Binding Factor), a protein of the fibrillar center (FC) and NPM1 (Nucleophosmin 1), a marker for the granular components (GC) (53,54) (Fig. 2A). However, in HeLa cells under hypoxia, there was significant reduction of the nucleolar signal of EXOSC10 with a concomitant increase in the nucleoplasmic signal (Fig. 2A and Suppl. Fig. 2) as documented by quantification (Fig. 2B). These data suggest that, in addition to de-SUMOylation, hypoxia also causes redistribution of EXOSC10 from the nucleolus to the nucleoplasm. Interestingly, hypoxia also caused similar redistribution to UBF and, to a lesser degree, NPM1 (Fig. 2A), suggesting that low oxygen conditions can cause notable restructuring of the overall nucleolar composition and architecture.

**Figure 2:**
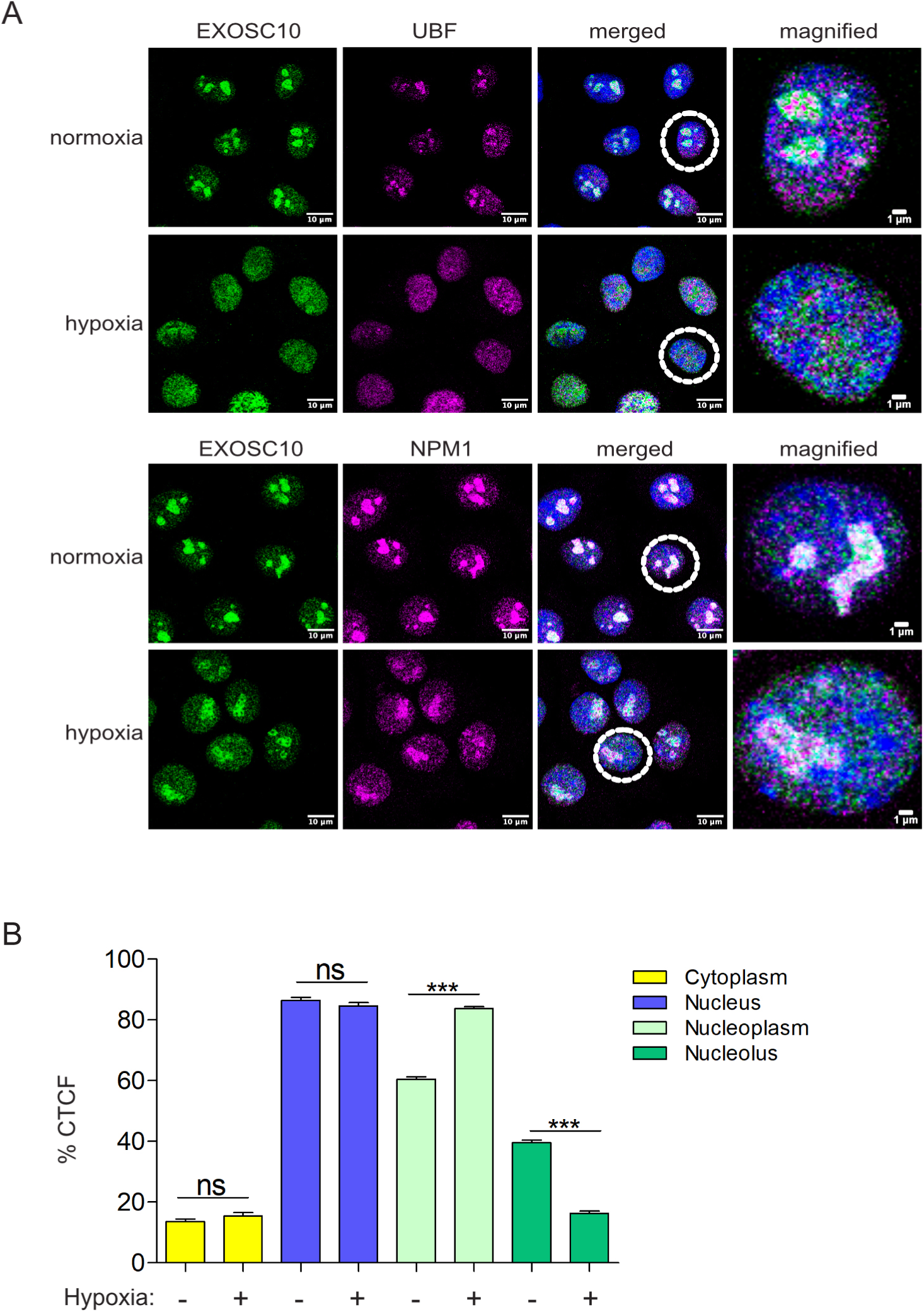
Hypoxia promotes redistribution of EXOSC10 from the nucleolus to the nucleoplasm. **A.** Indirect immunofluorescence (IF) analysis of HeLa cells incubated in normoxia or hypoxia for 24 h using antibodies against EXOSC10 and UBF, or EXOSC10 and NPM1 as indicated. DAPI was used for visualization of nuclear material. A cell, marked with dashed circle, is represented in magnified view as the last panel on the right for each case. Scale bar represent 10 μm or 1 μm as indicated. **B.** Graph representing quantification of the EXOSC10 IF signal (corrected cell fluorescence-CTCF) in cytoplasm, nucleus, nucleoli and nucleoplasm, from a total of 100 cells for each condition (three independent experiments) expressed as percentage of the total cell fluorescence, as mean ± standard error. For comparisons between two groups one-way ANOVA was used (***: P<0.001, n.s.: not significant).

### 3.3 SENP3 is the isopeptidase for SUMOylated EXOSC10 but does not mediate its hypoxia-induced deSUMOylation

As already mentioned, the hypoxia-induced reduction of EXOSC10 SUMOylation may be mediated by either inhibition of EXOSC10 SUMOylation or stimulation of EXOSC10 de-SUMOylation. To test the latter, we sought to identify the SUMO-isopeptidase responsible for de-conjugating SUMO from EXOSC10. SENPs represent the main family of SUMO-specific isopeptidases and two out of the six SENPs encoded in mammalians (55), SENP3 and SENP5, target many nucleolar proteins and are themselves enriched in the nucleolar compartment. SENP5 has been shown to have a clear preference for SUMO-2/3 substrates (56), so we first tested the involvement of SENP3 in modulating EXOSC10 SUMOylation. Silencing of SENP3 expression enhanced SUMOylation of EXOSC10 both under normoxia and hypoxia without, however, abolishing the difference in EXOSC10 SUMOylation between the two oxygen conditions (Fig. 3A). First, this suggests that SENP3 can release SUMO from SUMOylated EXOSC10. This is further supported by the physical association between EXOSC10 and SENP3 shown by their co-IP (Fig. 3B) as well as by the fact that silencing neither SENP1 nor SENP2 expression (two nuclear SENPs with most known substrates) affected EXOSC10 SUMOylation (Suppl. Fig. 3A). Second, it appears that SENP3 is not responsible for the reduction of EXOSC10 SUMOylation under hypoxia. In agreement with this scenario, expression of SENP3 was not increased by hypoxia (Fig. 3C) and hypoxic treatment reduced the association between EXOSC10 and SENP3 (Fig. 3B).

**Figure 3:**
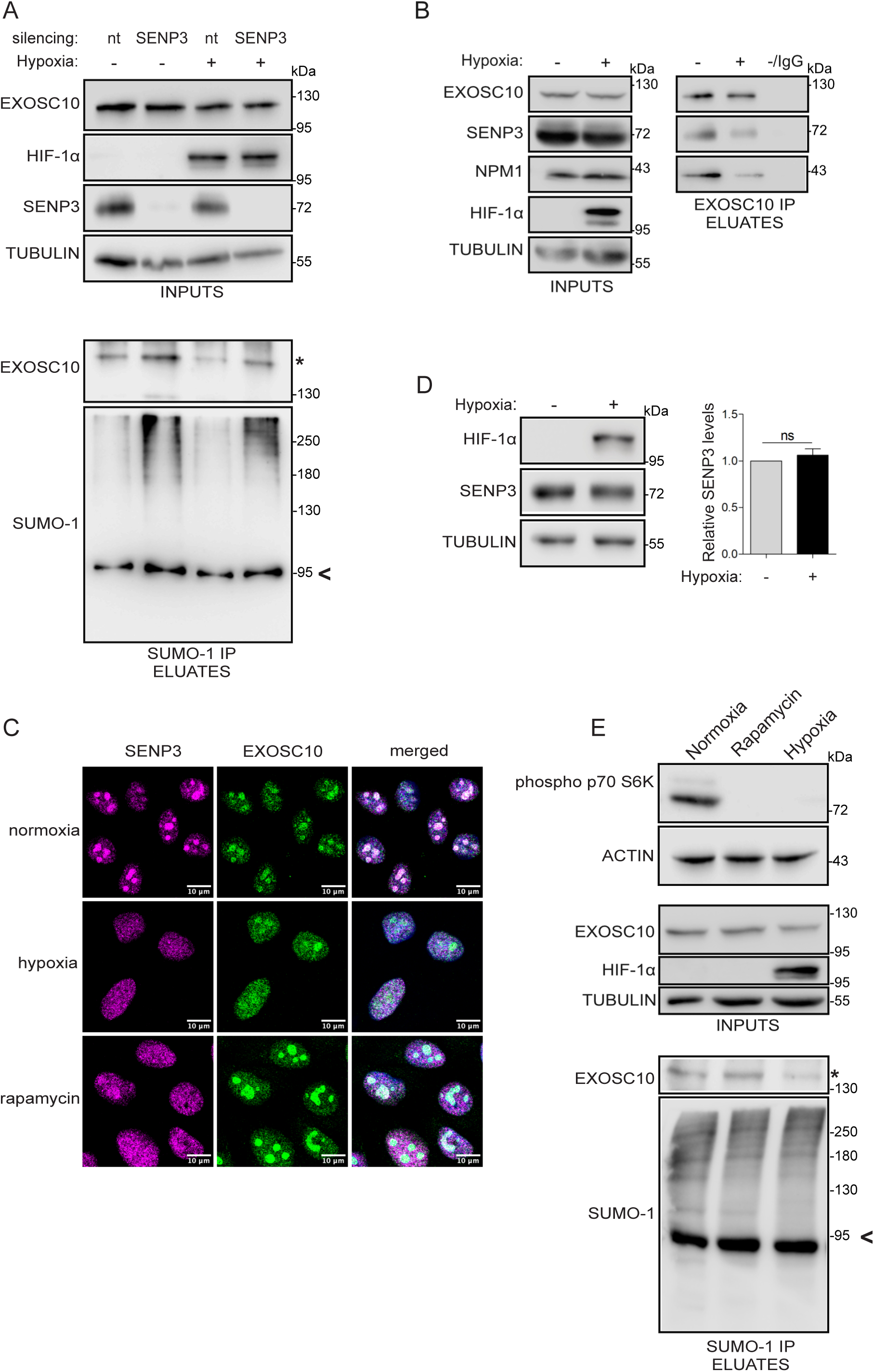
SENP3 is the isopeptidase for SUMOylated EXOSC10 but does not mediate its hypoxia-induced deSUMOylation. **A.** HeLa cells were transfected with a non-targeted siRNA (nt), or siRNAs against SENP3 and 24 h post-transfection, were incubated in normoxia (-) or hypoxia (+). Cell lysates were subjected to SUMO-1 IP. Inputs and eluates were analyzed by immunoblotting using the indicated antibodies. **B.** Cells were incubated in normoxia (-) and hypoxia (+) and lysates were subjected to EXOSC10-IP. An equivalent amount of lysate was also incubated with IgG-beads and served as negative IP control. Inputs and eluates were analyzed by immunoblotting using the indicated antibodies. **C.** Indirect immunofluorescence analysis of HeLa cells incubated in normoxic/hypoxic conditions as in (B) or treated with rapamycin for 6 h, using the indicated antibodies. DAPI was used for visualization of nuclear material. Scale bar represents 10 μm. **D.** HeLa cells were treated as in (B) and the endogenous expression levels of SENP3 were analyzed by immunoblotting. Beta-tubulin served as a loading control and HIF-1α was monitored to confirm the hypoxic response. Quantification of SENP3 protein levels (normalized to their tubulin signal) is shown on the right of the blot. Values are the mean of three independent experiments and are shown as fold increase compared to normoxic control (grey) and as mean ± standard error. For comparisons between groups unpaired t-test was used (n.s.: not significant). **E.** HeLa cells were treated as in (C), and cell lysates were subjected to SUMO-1 IP. Inputs and eluates were analyzed by immunoblotting using the indicated antibodies. The SUMOylated version of EXOSC10 is indicated with an asterisk (*). SUMOylated RanGAP1 is shown with an arrow and was indicated as a marker for equal precipitation by anti-SUMO1. Beta-tubulin was used as loading control.

To confirm these conclusions, the localization of SENP3 in normoxia and hypoxia was examined. Under normoxia, SENP3 co-localized with EXOSC10 predominantly in nucleoli, but under hypoxia, SENP3 redistributed from the nucleolus into the nucleoplasm, similar to EXOSC10 (Fig. 3D). This observation raised the possibility that, although not directly responsible for hypoxia-induced reduction in EXOSC10 SUMOylation, SENP3 could contribute to the relocalization and deSUMOylation of EXOSC10 under hypoxia by its own re distribution within the nucleus.

To test this possibility, we sought to trigger nuclear re-distribution of SENP3 through means other than hypoxic treatment. It has been previously shown that nucleolar targeting of SENP3 occurs via interaction with NPM1 and requires mTOR-mediated phosphorylation of SENP3 (60). Indeed, inhibition of mTOR by rapamycin, as confirmed by the reduction in p70S6K phosphorylation, which was also observed under hypoxia (Fig. 3E, upper panel), resulted in the release of SENP3 from the nucleolus to a similar extent as under hypoxia (Fig. 3D). However, upon rapamycin treatment EXOSC10 remained nucleolar (Fig. 3D) and its SUMOylation was not detectably affected (Fig. 3E, lower panel). Nucleolar release and dephosphorylation of SENP3 (Suppl. Fig 3B) under hypoxia can therefore be attributed to inhibition of the mTOR pathway, which is known to be affected by low oxygen (57), However, although SENP3 can deSUMOylate EXOSC10 under both normoxic and hypoxic conditions, dephosphorylation and/or translocation of SENP3 are not, at least solely, responsible for the nucleolar release of EXOSC10 and its deSUMOylation in hypoxia.

### 3.4 USP36 is the E3 SUMO ligase for EXOSC10 and hypoxia-induced disruption of their association inhibits EXOSC10 SUMOylation

As the loss of EXOSC10 SUMOylation under hypoxia cannot be explained by the effects of hypoxia on SENP3 phosphorylation and localization, we sought to identify the E3 SUMO ligase that catalyzes EXOSC10 SUMOylation, since its inhibition under hypoxia could rationalize the loss of EXOSC10 SUMOylation. To this end, the effects of silencing the expression of either of the two E3 SUMO ligases known to reside in the nucleolus, namely p14^ARF^ and USP36 (58,59) on EXOSC10 SUMOylation were monitored. Knocking down USP36, but not p14^ARF^ under normoxia markedly reduced modification of EXOSC10 by SUMO-1 to a similar extent as under hypoxia (Fig. 4A). Furthermore, USP36 could be detected in a protein complex together with EXOSC10 and NMP1 (Fig. 4B), strongly suggesting that USP36 can act as E3 SUMO ligase for EXOSC10. Although the protein expression level of USP36 was not affected by hypoxia (Fig. 4C), hypoxic treatment abolished the interaction between USP36 and EXOSC10 without significantly affecting the interaction between NPM1 and EXOSC10 (Fig. 4B). In addition, although under normoxia USP36 localized in the nucleolus, the co-localization with EXOSC10 in this compartment was largely lost in cells under hypoxia (Fig. 4D). Taken together, these results support the hypothesis that loss of EXOSC10 SUMOylation under hypoxia can be attributed to the physical separation of EXOSC10 from its E3 SUMO specific ligase, USP36, triggered by the nucleolar release of EXOSC10.

**Figure 4:**
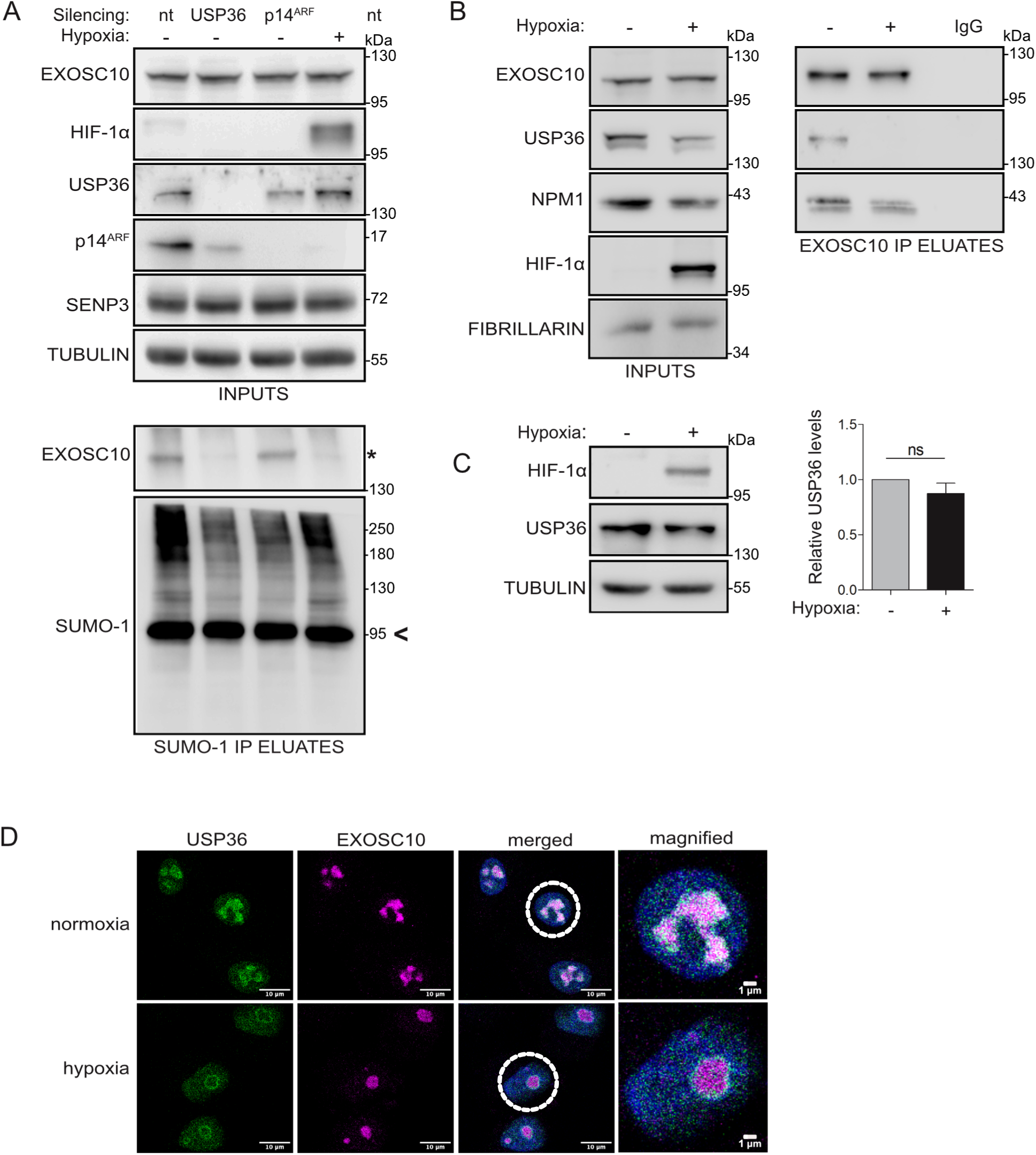
USP36 is the E3 SUMO ligase for EXOSC10 and the hypoxia-induced disruption of their association inhibits SUMOylation of EXOSC10. **A.** HeLa cells were transfected with non-targeted siRNA (nt), or siRNAs against USP36 or p14^ARF^ and 24 h post-transfection, were incubated in normoxia (-) or hypoxia (+). Cell lysates were subjected to SUMO-1 IP. Inputs and eluates were analyzed by immunoblotting using the indicated antibodies. **B.** Cells were incubated in normoxia (-) and hypoxia (+) and nuclear extracts were subjected to EXOSC10-IP. Equal amount of lysate was also loaded on IgG-beads and was used as negative IP control. Inputs and eluates were analyzed by immunoblotting using the indicated antibodies. **C.** HeLa cells were treated as in (B) and the endogenous expression levels of SENP3 were analyzed by immunoblotting. Beta-tubulin served as a loading control and HIF-1α was monitored to confirm the hypoxic response. Quantification of SENP3 protein levels (normalized to their tubulin signal) is shown on the right of the blot. Values are the mean of three independent experiments and are shown as fold increase compared to normoxic control (grey) and as mean ± standard error. For comparisons between groups unpaired t-test was used (n.s.: not significant). **D.** Indirect immunofluorescence analysis of HeLa cells incubated in normoxic/hypoxic conditions as in (B) using the indicated antibodies. DAPI was used for visualization of nuclear material. Scale bar represents 10 μm. The SUMOylated version of EXOSC10 is indicated with an asterisk (*). SUMOylated RanGAP1 is shown with an arrow and was indicated as a marker for equal precipitation by anti-SUMO1. Beta-tubulin was used as loading control.

### 3.5 K583 is the main site for modification of EXOSC10 by SUMO1

To explore the biological significance of EXOSC10 SUMOylation and its reduction under hypoxia, it was necessary to express a SUMO-deficient form of EXOSC10. Previous high-throughput SUMO-proteomic analysis identified at least 10 lysine residues in EXOSC10 as candidates for SUMOylation (60) and a triple lysine to arginine substitution (K168R, K201R and K583R) in EXOSC10 was observed to significantly reduce EXOSC10 SUMOylation (61). Two of these lysine residues, K168 and K583, reside in a classical SUMO consensus motif (ψKxE/D, Fig. 5A). To map the exact SUMOylation site(s) of EXOSC10, we individually substituted the lysine residues in positions 168, 201 and 583 (Fig. 5A) for arginines. Recombinant GST-fusion EXOSC10 proteins in wild-type form or carrying these amino acid substitutions (GST-EXOSC10 WT, K583R, K168R and K201R) were then purified from *E. coli* (Suppl. Fig 4A, left panel) and used as substrates for generic *in vitro* SUMOylation assays containing the E2 SUMO conjugating enzyme Ubc9, the catalytic fragment of E3 ligase RanBP2 (amino acids 2304-3062) and ATP. Reactions lacking ATP showed no SUMOylation and were used as negative controls. Reactions containing ATP led to SUMOylation of GST-EXOSC10 WT by UBC9 in a dose-dependent manner (Fig. 5B). With the optimal assay conditions, the K168R or K201R forms of EXOSC10 were SUMOylated but the K583R version lacked detectable SUMOylation, strongly suggesting that lysine 583 is the main SUMO modification site of EXOSC10.

**Figure 5:**
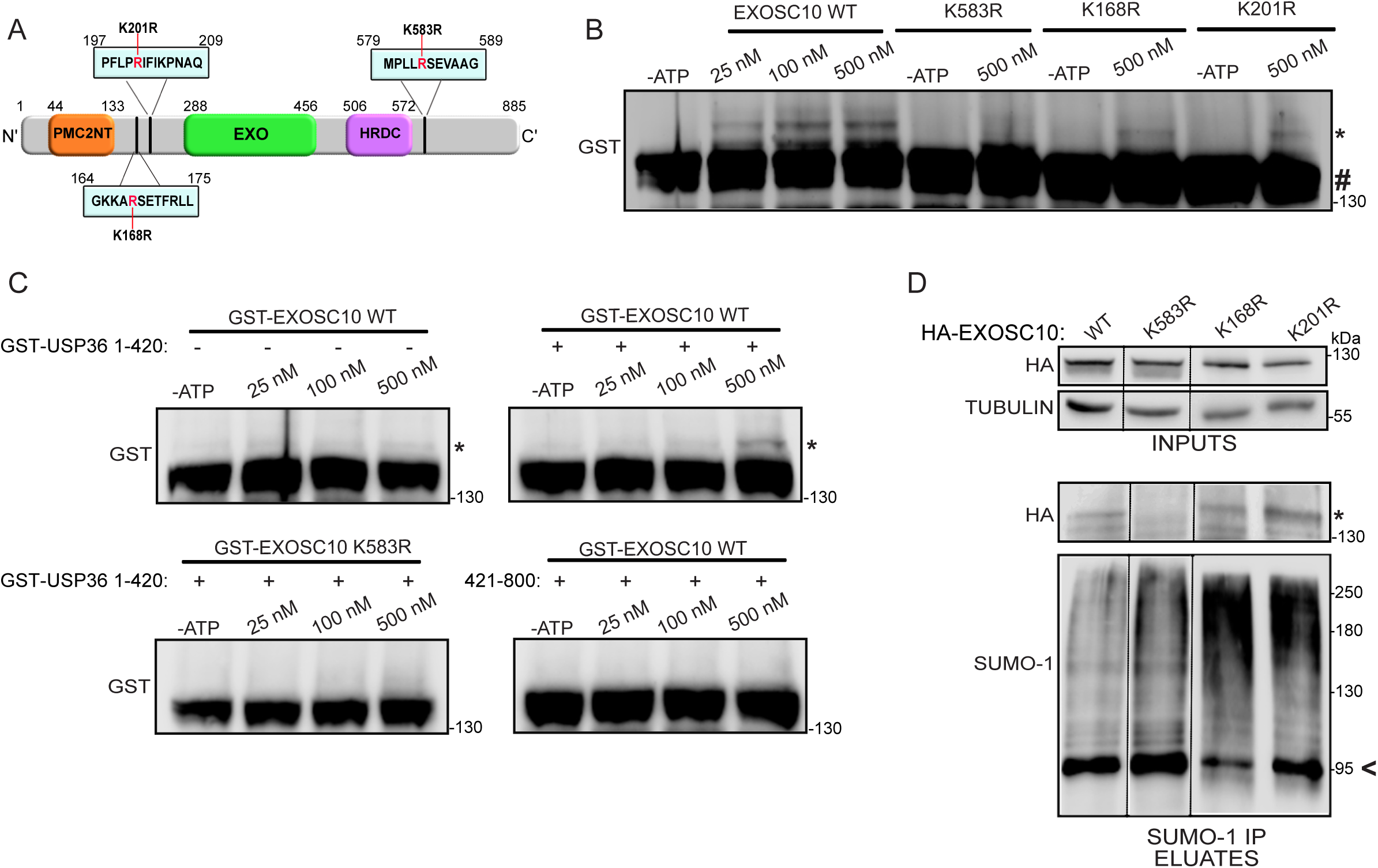
Lysine 583 is the main SUMOylation site of EXOSC10. **A.** Schematic diagram of EXOSC10. The positions of the polycystin 2 N-terminal (PMC2NT), exonuclease (EXO) and helicase and RNase D C-terminal (HRDC) domains are shown indicated. The positions of the SUMOylation motifs are marked and the amino acid substitutions used (K583R, K168R, K201R) are highlighted in red. **B.** *In vitro* SUMOylation reactions of the indicated GST-tagged recombinant EXOSC10 proteins (0,25 μM) incubated with 100 nM His-RanBP2-BD3-4 and increasing amounts of UBC9 (indicated), in the presence or absence of ATP. **C.** *In vitro* SUMOylation reactions of the indicated GST-tagged recombinant EXOSC10 proteins (0,25 μM) incubated with 100 nM GST-USP36 (1–420) or GST-USP36 (421–800) and increasing amounts of UBC9 (indicated), in the presence or absence of ATP. The reactions in (B) and (C) were further analyzed by immunoblotting using an antibody against GST. **D.** Hela cells expressing the indicated HA-EXOSC10 variants were subjected to SUMO-1 IP. Inputs and eluates were analyzed by immunoblotting using the indicated antibodies. In all cases, the SUMOylated version of EXOSC10 is indicated with an asterisk (*). SUMOylated RanGAP1 is shown with an arrow and was indicated as a marker for equal precipitation by anti-SUMO-1. Beta-tubulin was used as loading control.

To verify that the same residue is also the target of USP36, the amino-terminal part of USP36 (amino acids 1-420) that possesses SUMO E3 ligase activity and its non-catalytic carboxy-terminal part (amino acids 421-800) (29) were purified as recombinant proteins from *E. coli* (Suppl. Fig. 4A) and used in the *in vitro* SUMOylation assay replacing RanBP2. Inclusion of USP36(1–420), but not USP36(421–800), caused detectable SUMOylation of wild-type EXOSC10 while the K583R mutant form of EXOSC10 could not be modified under the same conditions (Fig. 5C). These *in vitro* data confirmed that USP36 is an E3 ligase for EXOSC10 and further show that it targets the lysine at position 583. This conclusion was further verified by over-expression of HA-tagged wild-type and K168R, K583R or K201R EXOSC10 in HeLa cells followed by SUMO-1 IP. All the EXOSC10 variants were similarly expressed, but while SUMOylation of the wild-type EXOSC10 and the K168R or K201R forms was readily detectable, SUMOylation of the K583R version was not observed (Fig. 5D), demonstrating that lysine 583 is the main SUMO modification site of EXOSC10 *in vivo*.

Before using the K583R variant as a SUMO-deficient form of EXOSC10 it was important to verify that the K583R substitution did not affect the catalytic activity of EXOSC10 by perturbing its overall structure. Bacterially expressed, unmodified recombinant GST-EXOSC10 WT and GST-EXOSC10 K583R, as well as the catalytically inactive GST-EXOSC10 D313A form (38), were therefore tested in an *in vitro* exoribonuclease assay using a synthetic RNA substrate (U_32_). The wild-type and the K583R mutant forms displayed similar enzymatic activities, in contrast to the inactive D313A form (Suppl. Fig. 4B), implying that the K583R mutation does not disturb the catalytic function of EXOSC10.

### 3.6 SUMO modification of EXOSC10 affects the expression of hypoxia-regulated transcripts

To construct a cell line rescue system to analyse phenotypes associated with expressing SUMO-deficient EXOSC10 K583R, we used the Flp-In T-REx system (20). Specifically, stably transfected HeLa Flp-In cell lines in which the expression of siRNA-resistant Flag-tagged EXOSC10 WT or EXOSC10 K583R could be induced by addition of tetracycline while expression of endogenous EXOSC10 was simultaneously knocked-down by siRNA-mediated silencing, were generated. A HeLa FlpIn cell line for the inducible expression of only the Flag-tag served as a control cell line. As shown in Fig. 6A, upon induction and siRNA treatment, these cells expressed Flag-EXOSC10 WT or Flag-EXOSC10 K583R at similar levels as well as endogenous EXOSC10 in unsilenced cells.

**Figure 6:**
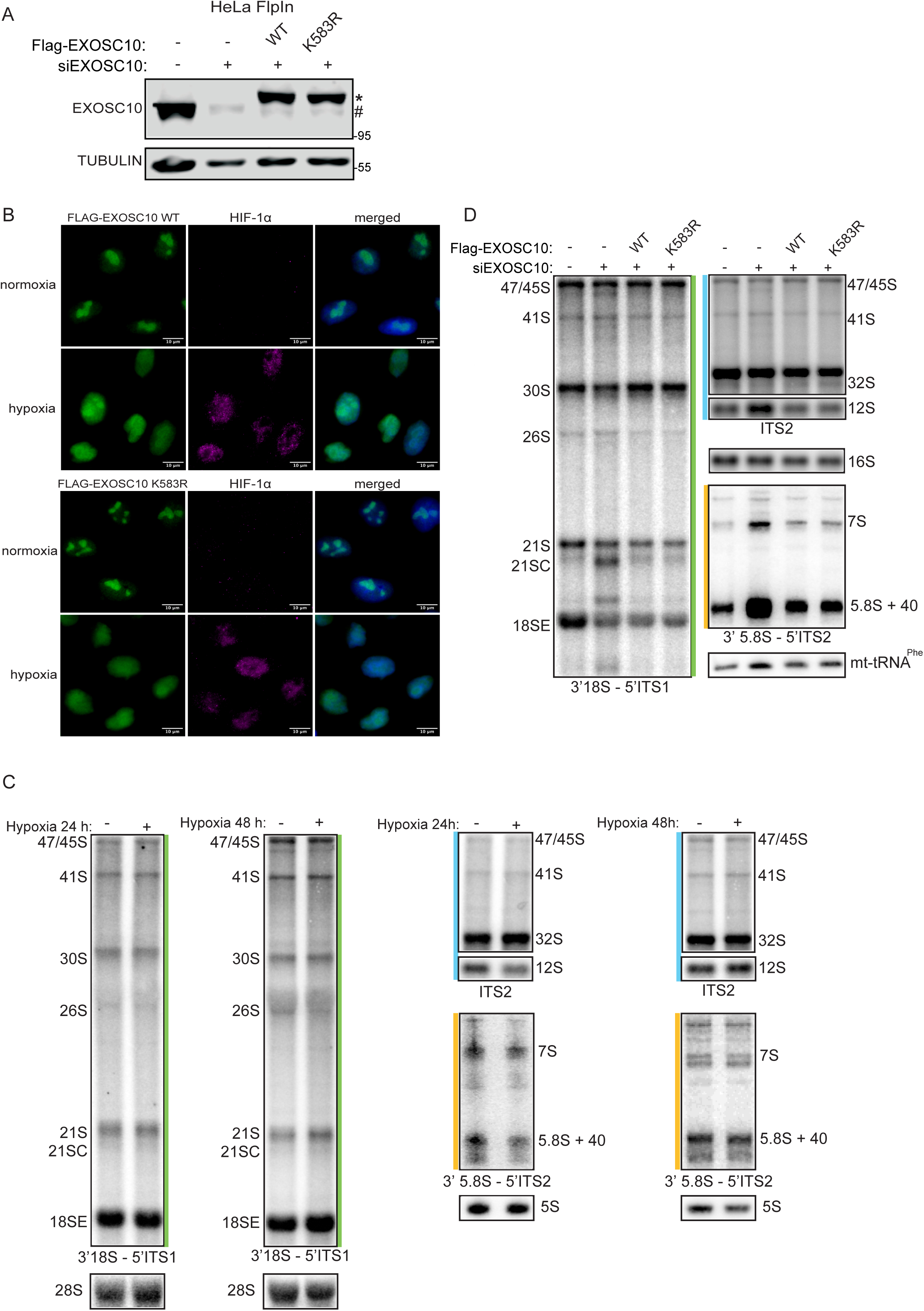
Pre-rRNA processing is not significantly affected by expression of non-SUMOylated EXOSC10 K583R or hypoxia treatment. **A.** Immunoblotting analysis using the indicated antibodies, of stably transfected HeLa FlpIn cell lines expressing either the 2xFLAG-His_6_ (FLAG) tag (-), or FLAG-EXOSC10 WT or FLAG-EXOSC10 K583R, in the presence of non-targeted siRNA (-) or a pool of siRNAs against EXOSC10. Exogenously expressed proteins are shown with an asterisk and endogenous EXOSC10 is shown with a hash mark (#). Beta-tubulin was used as loading control and MW are shown on the right. **B.** Stably transfected HeLa FlpIn cells expressing either FLAG-EXOSC10 WT or FLAG-EXOSC10 K583R were grown on coverslips for 24 h in normoxia or hypoxia and subjected to indirect immunofluorescence analysis using antibodies against the FLAG-tag and HIF-1α as control for the hypoxic conditions. DAPI was used for visualization of nuclear material. Scale bar represents 10 μm. **C.** RNAs extracted from stably transfected HeLa FlpIn cell lines expressing either the FLAG tag (-), or FLAG-EXOSC10 WT or FLAG-EXOSC10 K583R, in the presence of non-targeted siRNA (-) or a pool of siRNAs against EXOSC10 were analyzed by northern blotting using probes complementary to the regions of the pre-rRNA indicated in Suppl. Fig. 5A. RNA loading was monitored using the mitochondrial (mt)16S rRNA and mt-tRNA^Phe^. **D.** HeLa cells were incubated in normoxia or hypoxia for 24 and 48h and pre-rRNAs were further analyzed by northern blotting as in (C).

Under normoxia, both stably expressed WT and SUMO-deficient Flag-EXOSC10 were nuclear and enriched in nucleoli (Fig. 6B, upper panels), i.e. exhibited the same localization as endogenous EXOSC10 (see Fig. 2). Under hypoxia, the nucleolar signals of both forms were weakened with concomitant increases of their signals in the nucleoplasm (Fig. 6B, lower panels), again analogous to endogenous EXOSC10 (Fig. 2). These data suggest that the SUMOylation status of EXOSC10 does not influence its steady-state nucleolar localization or its hypoxia-induced nucleolar release. Redistribution of EXOSC10 to the nucleoplasm under hypoxia is, therefore, likely not caused by deSUMOylation but may instead be the reason for the reduced SUMOylation.

EXOSC10 has a well-characterized role in the maturation of rRNA precursors. To explore if hypoxia or the SUMOylation status of EXOSC10 affects the levels of pre-rRNA intermediates, pre-rRNA processing was analyzed in the cells exposed to hypoxia as well as those depleted of endogenous EXOSC10 alone or complemented with expression of either wild-type or SUMO-deficient (K583R) EXOSC10. Northern blotting using probes to detect all the major pre-rRNA intermediates (Supp. Fig. 5A) showed that the levels of rRNA precursors isolated from HeLa cells grown under normoxia or hypoxia for 24 or 48 h did not significantly differ, implying little or no effect of hypoxia on pre-rRNA processing (Fig. 6C and Suppl. Fig. 5B). Upon silencing of endogenous EXOSC10, accumulation of several pre-rRNA species, e.g. 21SC, 7S and 5.8S+40 (Fig. 6D and Suppl. Fig. 5C) was observed, consistent with the known roles of EXOSC10 in ribosome biogenesis (38,39). Induction of either FLAG-EXOSC10 WT or FLAG-EXOSC10 K583R in cells depleted of endogenous EXOSC10 rescued the defective phenotype and the levels of these pre-rRNAs were restored to normal in both cases (Fig. 6D and Suppl. Fig. 5C). These findings suggest that neither hypoxia and the associated nuclear re-distribution of EXOSC10 nor loss of EXOSC10 SUMOylation severely affect the processing of rRNA precursors.

Beyond rRNA precursors, EXOSC10 has many other targets so to then analyse the involvement of EXOSC10 SUMOylation and its regulation by hypoxia on the transcriptome, we utilized RNA-seq. Total RNAs were extracted from HeLa cells grown under normoxia or hypoxia, as well as those expressing either the WT or K583R version of EXOSC10. Following rRNA depletion, RNAs were fragmented and converted to a cDNA library that was analyzed by next-generation sequencing. Principal component analysis of the obtained sequencing data showed good reproducibility between replicate datasets and clear distinctions between samples derived from cells under normoxia and hypoxia, as well as those from cells expressing WT or K583R versions of EXOSC10 (Fig. 7A).

**Figure 7:**
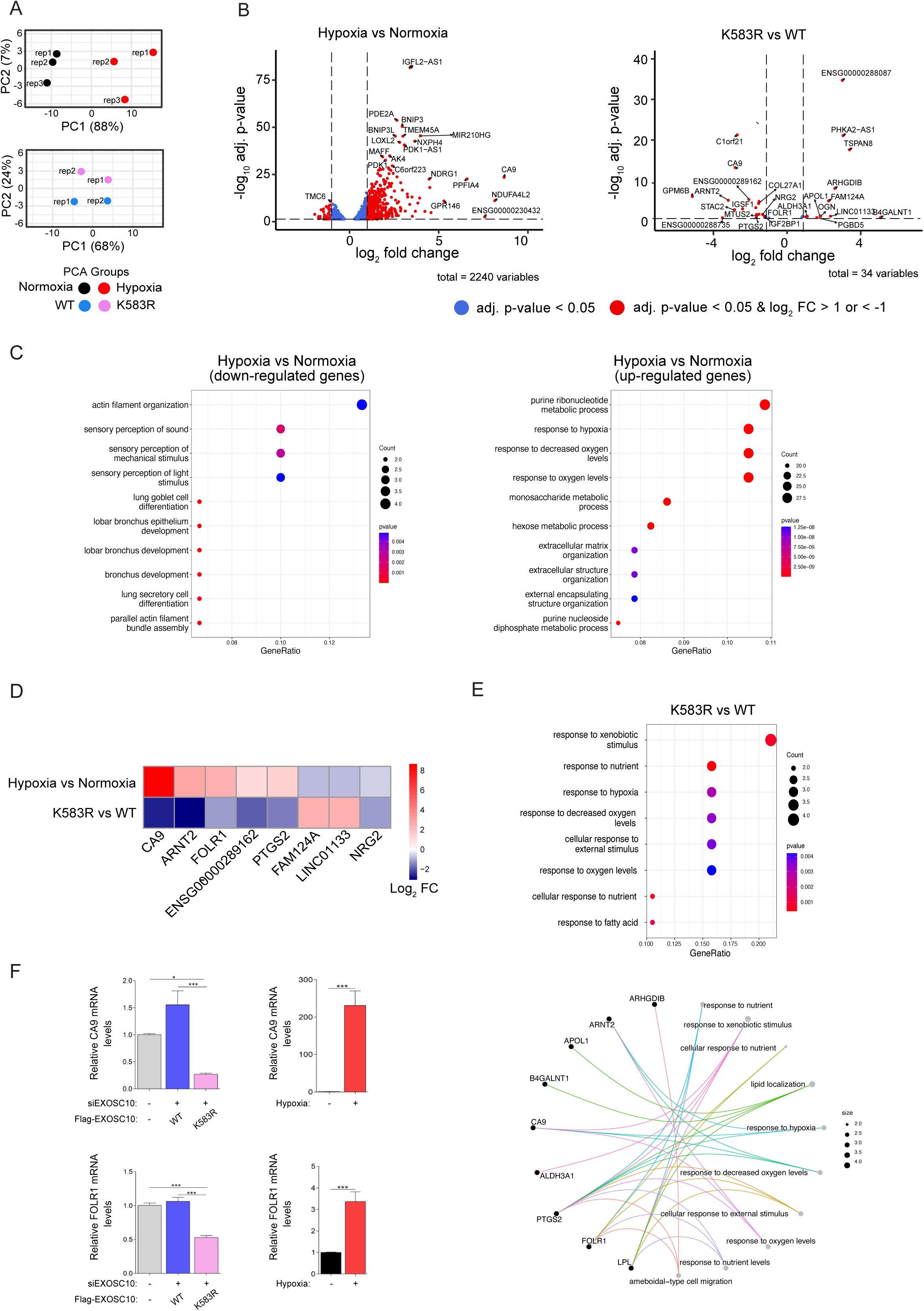
Hypoxia-mediated deSUMOylation of EXOSC10 affects the transcriptome profile. **A-D.** Total RNA extracted from HeLa FlpIn cells after 24 h of hypoxia or normoxia treatment, and from stably transfected HeLa FlpIn cell lines expressing either FLAG-EXOSC10 WT (WT) or FLAG-EXOSC10 K583R (K583R) depleted of endogenous EXOSC10 (siEXOSC10) were subjected to RNA-seq. **A.** Principal component analysis (PCA) for the compared groups hypoxia *versus* normoxia and K583R *versus* WT. **B.** Volcano plots showing genes with significant differential expression (adjusted p-value < 0.05) when comparing the conditions hypoxia *versus* normoxia and K583R *versus* WT. DESeq2 was used to normalize the read counts and to calculate the differential expression of genes between two conditions. Red dots indicate up-regulated genes (log_2_ FC > 1) or down-regulated genes (log_2_ FC < -1). Vertical dashed lines indicate the fold-change cut-offs log_2_ FC > 1 and log_2_ FC < -1. Horizontal dashed line indicates the threshold of adjusted p-value (adj. p-value < 0.05). **C.** Dotplots highlighting Top10 GO-pathways enriched in HeLa cells using the DE gene lists generated by group comparisons (left: down-regulated genes of hypoxia *versus* normoxia comparison (log_2_ FC < -1, p adj < 0.05), right: up-regulated genes of hypoxia *versus* normoxia comparison, (log2FC > 1, p adj < 0.05). **D**. Heatmap indicating the log2 FC values of selected genes with significant differential expression levels between hypoxia *versus* normoxia and K583R *versus* WT. Shown genes have log_2_ FC > 1 or log_2_ FC < -1 in K583R *versus* WT. Color spectrum from red to blue represent expression of up-and down-regulated genes, respectively. **E.** Dotplot (up) highlighting Top8 GO-pathways enriched and Cnet plot (down) highlighting Top10 GO-pathways enriched with associated genes in HeLa FlpIn cells expressing the FLAG-EXOSC10 K583R compared with cells expressing the FLAG-EXOSC10 WT (log2FC > 1/ < -1, adj. p-value < 0.05). Size of the dots refer to the number of genes enriched with the GO-term. **E.** CA9 and FOLR1 expression levels were determined by RT-PCR as indicated. Results are shown as fold increase (Mean SEM) in relation to the control conditions (-/-) and represent the mean of two for the rescue system and three for the normoxic/hypoxic conditions biological independent experiments performed in technical duplicates (n = 4/6, ****P<0.0001, ***P < 0.001, **P<0.01, *P<0.05, ns: not-significant; Statistical variance between two groups of values was calculated by one-way ANOVA).

When compared to normoxia, hypoxia significantly altered the mRNA levels of 475 genes (427 upregulated and 48 downregulated, p-adj<0.05 and log_2_FC >1 or < - 1, Fig. 7B and Suppl. Excel File 1). As expected, the set of upregulated genes was enriched for genes involved in the response to oxygen levels and various metabolic processes, while there was enrichment of genes involved in actin organization and sensory perception in the downregulated set (Fig. 7C). Expression of the EXOSC10 K583R SUMO-deficient mutant in cells depleted of endogenous EXOSC10 compared to the expression of the WT form, affected the expression of 31 genes (19 downregulated and 12 upregulated), most of which are involved in the cellular response to environmental stimuli such as oxygen, nutrients and xenobiotics (Fig. 7D and E), supporting that regulation of EXOSC10 SUMOylation is important for the hypoxic response.

Importantly, several of the mRNAs affected by expression of the EXOSC10 K583R SUMO-deficient mutant were also affected by hypoxia (Fig. 7D). For example, the mRNAs encoding CA9, ARNT2, FOLR1, PTGS2 were upregulated under hypoxia (compared to normoxia) and downregulated upon EXOSC10 K583R expression (compared to EXOSC10 WT overexpression), implicating EXOSC10 deSUMOylation in their regulation by hypoxia. The RNA-seq results were validated by RT-PCR (Fig. 7F) for two representative mRNAs: CA9, a well-known hypoxia-inducible gene, coding for carbonic anhydrase 9, and FOLR1, coding for folate receptor 1. Consistent with the RNA-seq data, both targets were strongly induced by hypoxia and down-regulated upon K583R expression.

## 3. Discussion

Activation of gene transcription mediated by HIFs plays a central role in the response to hypoxia (7,62). However, in recent years, non-transcriptional processes, such as protein synthesis (57,63) and mRNA stability, have also emerged as adaptive mechanisms to low oxygen conditions (15,31). Examples include the inhibition of nonsense-mediated RNA decay (NMD) by anoxia (<0.1% O_2_) via an eIF2α phosphorylation mechanism (64) and hypoxia-inducible stabilization of mRNAs by binding of the HuR protein to AU rich sequences in the 3’UTR regions (65). More recently, it was also shown that hypoxia reduces the overall mRNA median half-life and the cellular mRNA and total RNA content in endothelial cells (66). Our study now provides evidence of crosstalk between the response to hypoxia and the RNA degradation machinery by showing that hypoxia leads to deSUMOylation of EXOSC10 in a HIF-independent manner. Furthermore, we have identified the enzymes responsible for the reversible SUMOylation of EXOSC10 and analyzed how hypoxia affects their expression and subcellular localization. Our data show that hypoxia causes translocation of EXOSC10 from the nucleolus to the nucleoplasm and dissociation from its nucleolar E3 ligase complex, explaining its lack of SUMOylation. Moreover, loss of EXOSC10 SUMOylation affects the expression level of hypoxia-relevant transcripts in a way that may offer advantages for the cellular adaptation to low oxygen conditions.

Previously, we and others (33) (49, 53, 61) identified EXOSC10 as a SUMOylated protein in large-scale proteomic screens. Interestingly, heat shock modulates SUMOylation of EXOSC10 (67) while cooling-induced modification of EXOSC10 by SUMO was shown to reduce its protein levels (62), suggesting that regulation of EXOSC10 SUMOylation may be a common strategy for fine tuning responses to different stress conditions. However, until recently the enzymes involved in EXOSC10 SUMOylation had not been characterized. Our data suggest that SENP3, as a SUMO-isopeptidase, and USP36, as an E3 SUMO ligase, mediate the reversible (de)SUMOylation of EXOSC10. The latter finding is in agreement with a study published during the preparation of this manuscript, which identified EXOSC10 as a USP36-interacting protein and target (68). We have further shown that SENP3 and USP36 reside in the same nucleolar complex with EXOSC10 and NPM1 in cells under normoxia. SENP3 is known to be the enzyme responsible for the deSUMOylation of NPM1(69) and USP36 is known to be involved in the SUMOylation of many nucleolar proteins such as the small nucleolar ribonucleoprotein (snoRNP) components Nop58 and Nhp2 (28). It is, therefore, likely that NPM1 creates a scaffold that recruits EXOSC10 in a nucleolar complex in which USP36 and SENP3 mediate the balance between SUMOylated and deSUMOylated forms of EXOSC10. This balance in disturbed under hypoxia, as, according to our data, exposure to low oxygen triggers changes in nucleolar protein composition by promoting the redistribution of EXOSC10, as well as UBF and SENP3 but not USP36, from the nucleolus to the nucleoplasm.

It has been previously shown that inhibition of the mTOR pathway causes mislocalization of SENP3 to nucleoplasm via its de-phosphorylation and detachment from NPM1 (70). We confirmed that treatment with rapamycin, an inhibitor of mTOR, under normoxia induced SENP3 to mislocalize, but strikingly, the localization and SUMOylation EXOSC10 were not affected. This strongly suggests that the effects of hypoxia on EXOSC10 localization and SUMOylation are not mediated by the dephosphorylation/mislocalization of SENP3 resulting from mTOR downregulation under low oxygen conditions. We, therefore, suggest that the changes in EXOSC10 localization and SUMOylation under hypoxia are a consequence of its loss of interaction with its SUMO E3 ligase USP36. We cannot, however, exclude that after the dissociation of EXOSC10 from the USP36-containing nucleolar complex and its translocation in the nucleoplasm, EXOSC10 may interact with other nuclear SUMO isopeptidases, which could shift the SUMO conjugation equilibrium towards deSUMOylation. In any case, the trigger for the hypoxia-induced displacement of EXOSC10 from the nucleolus still remains unknown. Early non-transcriptional hypoxic events such as intracellular formation of ROS or activation of kinase pathways could influence the overall structural organization of the nucleolus and the interactions involving EXOSC10.

An interesting question raised by our findings is what the effects of SUMOylation on the functions of EXOSC10 are and whether they are important for cellular adaptation to hypoxia. A previous study in HEK293 cells showed that exposure to low temperature (cooling) increased global SUMOylation of proteins as well as SUMOylation of EXOSC10, which caused suppression of EXOSC10 expression and 3′ pre-rRNA processing defects (61). By contrast, our data show that hypoxia-induced deSUMOylation does not affect the level of EXOSC10 protein in HEK293 or HeLa cells. Furthermore, our experiments demonstrate that the EXOSC10 K583R version lacking the SUMOylation site was as efficient as the wild-type form of EXOSC10 in complementing the pre-rRNA processing defects caused by silencing the expression of endogenous EXOSC10. This last finding is in contrast with the very recent study identifying EXOSC10 as a target for USP36, which suggested that deSUMOylation of EXOSC10 mildly impairs maturation of the 5.8S rRNA (68). It remains unclear why a similar effect is not observed when non-SUMOylatable Flag-tagged EXOSC10 K583R is transiently expressed to a similar level as endogenous EXOSC10 as re-expression of an equivalent wild-type EXOSC10 fully rescues the pre-rRNA processing defects caused by lack of EXOSC10. Notably, no significant pre-rRNA processing defects were observed in cells exposed to hypoxia, despite the fact that under the same conditions EXOSC10 almost completely loses its SUMOylation. The lack of effect of hypoxia on pre-rRNA processing is in line with a previous study showing that incubation of cancer cells under hypoxia combined with acidosis causes a VHL-dependent reduction in rDNA transcription and defects in ribosome biogenesis, but that hypoxia alone does not induce such a phenotype (12). Therefore, the extent of EXOSC10 deSUMOylation caused by hypoxia apparently does not cause a functionally significant defect in pre-rRNA maturation.

EXOSC10 is a multifunctional ribonuclease that also plays a role in the maturation/degradation of other RNA types. Interestingly, we identified several genes, the expression of which was altered in cells expressing the SUMO-deficient form of EXOSC10 (K583R) instead of the wild-type protein. A subset of these genes was also dysregulated by hypoxia, showing that the hypoxia-induced alteration of EXOSC10 SUMOylation has functional significance as it determines the level of transcripts coding for proteins required for adaptation to low oxygen conditions. In the case of the *CA9* and the *FOLR1* mRNA, their expression is suppressed by deSUMOylation of EXOSC10. This type of regulation might be important as part of a homeostatic feedback loop, triggered by deSUMOylation of EXOSC10 that reduces levels of hypoxia-dependent transcripts, such as CA9 and FOLR1, during prolonged hypoxia and can be a useful strategy for fine tuning responses to low oxygen conditions at the post-transcriptional stage.

Another question raised by our findings, is how the SUMOylation dynamics of EXOSC10 affect positively and negatively the fate of different mRNA transcripts. It is possible that SUMOylation of EXOSC10 affects its binding to RNA. As a recent example in favour of this hypothesis, SUMOylation of the RNA binding factor YTH domain family 2 (YTHDF2) increased its binding to and promoted the degradation of m^6^A-containing mRNAs (71). Interestingly, SUMOylation of YTHDF2 was increased under hypoxia and was reduced under oxidative stress. EXOSC10 is modified by SUMO-1 at K583, a residue present at the end of the helicase and RNase D carboxy terminal (HRDC) domain, which is known to regulate the activity of the catalytic exoribonuclease domain (EXO)(72). It is also close to a domain (amino acids 741-885) that in Rrp6, the yeast orthologue of EXOSC10, is called the ‘lasso’ and is used for recruiting RNAs to promote their decay (73). It is also possible that SUMOylation of K583 leads to conformational changes affecting the interaction of EXOSC10 with RNA-binding factors. The N-terminal domain of EXOSC10, PMC2NT, is known to bind to exosome cofactor C1D providing a scaffold for interaction with the RNA helicase MTR4 and exosome cofactor MPP6 (74,75), two proteins that facilitate degradation of poly(A)+ substrates by EXOSC10 in association with the RNA exosome (76). MTR4 is part of different nuclear RNA degradation complexes, the Trf4/5-Air1/2-Mtr4 polyadenylation (TRAMP) (77), the NEXT (nuclear exosome targeting) and the PAXT (poly(A) tail exosome targeting) complex, that direct the RNA exosome-mediated degradation of several different classes of RNA (76,78). *In silico* analysis (https://sumo.biocuckoo.cn) predicts the presence of several putative SUMO-interacting motifs (SIM) in the MTR4 sequence (79), which could potentially mediate the interaction between MTR4 and the SUMOylated form of EXOSC10 in the context of distinct RNA degradation complexes.

In summary, this study provides new information on the crosstalk between the response to hypoxia and the RNA degradation machinery. Although hypoxia has been known to affect mRNA stability (64) (65) (66), deciphering the underlying mechanisms has been challenging. Here, we reveal hypoxia-driven nucleolar changes that lead to deSUMOylation of EXOSC10 and subsequent differential expression levels of mRNA transcripts of hypoxia-inducible genes. This fine-tuning of gene expression via post-translational regulation of RNA degradation may be important for optimal cellular response and adaptation to low oxygen conditions.

## Data availability

The RNA-seq datasets for HeLa cells grown in normoxia and hypoxia, and those for cells depleted of endogenous EXOSC10 and ectopically expressing EXOSC10 WT or K583R are deposited in Gene Expression Omnibus (GEO) database [http://www.ncbi.nlm.nih.gov/geo/] under the accession code GSE229894. RNA-seq data are currently under quarantine and will be released upon publication in a peer reviewed journal.

## Supplementary data

Supplementary data are available in a different folder (Supporting info)

## Author Contributions

C.F., C.T. and G.C. performed experimental research, G.C., K.E.B. and G.S. designed research, G.C. and K.E.B. supervised experiments, G.C., G.S. and K.E.B. contributed reagents/analytic tools; S.P., C.T. and E.N performed bioinformatic analyses, E.N. supervised bioinformatic analysis, G.C. wrote the paper; C.F., G.S, E.N. and K.E.B. helped writing the paper.

## Acknowledgments

We thank Prof. Dr. Mushui Dai (OHSU, Oregon, US) for kindly providing USP36 plasmids. We acknowledge infrastructure support from the project “SyntheticBiology: From omics technologies to genomic engineering (OMIC-ENGINE) (MIS 5002636) which is implemented under the action “reinforcement of the Research and Innovation Infrastructure” funded by the Operational Programme “Competitiveness, Entrepreneurship and Innovation (NSRF 2014-2020) and co-financed by Greece and the European Union (European Regional Development Fund) and Dr. A.M. Psarra and I. Tsaltas (University of Thessaly, Greece) for their technical help. We thank Prof. Dr. Dörthe M. Katschinski (University of Göttingen, Germany) for the use of their hypoxia working station.

