## Supplementary data for "Hypoxia-driven deSUMOylation of EXOSC10 promotes adaptive changes in the transcriptome profile"

**Supplementary Table S1:** List of the DNA oligonucleotides used in this study.

| oligonucleotides | Sequence | Application | Reference |
|---|---|---|---|
| EXOSC10 For BamHI | 5'-TCAGGATCCATGGCGCCAC-3' | Molecular cloning pcDNA 3.1-HA-EXOSC10/ pBluescript II SK (+/-)-EXOSC10 | Sigma-Aldrich, StLouis, MO, USA |
| EXOSC10 Rev XbaI | 5'-CTCTAGATCATCTCTGTGGCC-3' | Molecular cloning pcDNA 3.1-HA-EXOSC10/ pBluescript II SK (+/-)-EXOSC10 | Sigma-Aldrich, StLouis, MO, USA |
| EXOSC10 K583R mutag. | 5'-AGATGCCCCTGCTCAGGTCTGAAGTTGCAG-3' | Site-directed mutagenesis ($EXOSC10_{K583R}$) | Eurofins |
| EXOSC10 K583R compl. mutag. | 5'-CTGCAACTTCAGACCTGAGCAGGGGCATCT -3' | Site-directed mutagenesis ($EXOSC10_{K583R}$) | Eurofins |
| EXOSC10 K168R mutag. | 5'TGGCAAAAAAGCAAGATCTGAAACTTTCCG-3' | Site-directed mutagenesis ($EXOSC10_{K168R}$) | Eurofins |
| EXOSC10 K168R compl mutag. | 5'-CGGAAAGTTTCAGATCTTGCTTTTTTGCCA-3' | Site-directed mutagenesis ($EXOSC10_{K168R}$) | Eurofins |
| EXOSC10 K201R mutag. | 5'-CACCATTTCTTCCTAGAATCTTCATCAAACC-3' | Site-directed mutagenesis ($EXOSC10_{K201R}$) | Eurofins |
| EXOSC10 K201R compl mutag. | 5'-GGTTTGATGAAGATTCTAGGAAGAAATGGTG-3' | Site-directed mutagenesis ($EXOSC10_{K201R}$) | Eurofins |
| EXOSC10 *D313A mutag.* | 5'-GAATTGTCAGGAATTTGCAGTTAACTTGGAGCACCACTCTTACAGG-3' | Site-directed mutagenesis ($EXOSC10_{D313A}$) | Eurofins |
| EXOSC10 D313A *compl mutag.* | 5'-CCTGTAAGAGTGGTGCTCCAAGTTAACTGCAAATTCTGACAATTC-3' | Site-directed mutagenesis ($EXOSC10_{D313A}$) | Eurofins |

| | | | |
|---|---|---|---|
| 3’18S – 5’ITS1 | 5’-CCTCGCCCTCCGGGCTCCGTTAATGATC-3’ | Northern blot probe | [1] |
| ITS2 | 5’-GCTCTCTCTTTCCCTCTCCGTCTTCC-3’ | Northern blot probe | Eurofins |
| 3’5.8S – 5’ITS2 | 5’GGGGCGATTGATCGGCAAGCGACGCTC-3’ | Northern blot probe | [2] |
| 16S | 5’-TTCTATAGGGTGATAGATTGGTCC-3’ | Northern blot probe | Eurofins |
| mt-tRNA$^{Phe}$ | 5’-TTGCTTTGAGGAGGTAAGCTACATAA-3’ | Northern blot probe | Eurofins |
| 28S | 5’- TACCTCTTAACGGTTTCACGCCCTCTTGAACTCTC-3’ | Northern blot probe | Eurofins |
| 5S | 5’-GAGATCAGACGAGATCGGGCGCGTTC-3’ | Northern blot probe | Eurofins |

**Supplementary Table S2:** List of plasmids used in this study.

| Plasmid | Application |
|---|---|
| pcDNA 3.1-HA-EXOSC10 WT | Overexpression in human cell lines |
| pcDNA 3.1-HA-EXOSC10 K583R | Overexpression in human cell lines |
| pcDNA 3.1-HA-EXOSC10 K168R | Overexpression in human cell lines |
| pcDNA 3.1-HA-EXOSC10 K201R | Overexpression in human cell lines |
| pcDNA5-2xFlag-His6-siinsEXOSC10 WT | Generation of stable HeLa FlpIn cell lines |
| pcDNA5-2xFlag-His6-siinsEXOSC10 K583R | Generation of stable HeLa FlpIn cell lines |
| pGEX-4T1-EXOSC10 WT | Protein expression in *E.coli* |
| pGEX-4T1-EXOSC10 K583R | Protein expression in *E.coli* |
| pGEX-4T1-EXOSC10 K168R | Protein expression in *E.coli* |
| pGEX-4T1-EXOSC10 K201R | Protein expression in *E.coli* |
| pGEX-4T1-EXOSC10 D313A | Protein expression in *E.coli* |
| pGEX-4T1-USP36 1-420 | Protein expression in *E.coli* (a kind offer from Dr. M.Dai) |
| pGEX-4T1-USP36 421-800 | Protein expression in *E.coli* (a kind offer from Dr. M.Dai) |

**Supplementary Table S3:** List of siRNAs used in this study.

| oligonucleotides | Sequence | Reference |
|---|---|---|
| AllStars Negative Control siRNA | Proprietary | Qiagen,Venlo,Netherlands |
| negative control siRNA (targeting firefly luciferase) | 5’- CGUACGCGGAAUACUUCGATT-3’ | [3] |
| siRNA against EXOSC10 #1 | 5’-GAAGGCAGCUGAGCAAACATT-3’ | [1] |
| siRNA against EXOSC10 #2 | 5’- UGAGCAGAGUAAUGCAGUATT-3’ | [1] |
| siRNA against EXOSC10 #3 | 5’- AGAUGAAAGUUACGGAUAUTT -3’ | [1] |
| siRNA against SENP3 | 5’- CAAUAAGGAGCUACUGCUATT -3’ | [4] |
| siRNA against USP36 Hs_USP36_3 (SI00138082) | 5’-CAAGAGCGTCTCGGACACCTA-3’ | Qiagen,Venlo,Netherlands |
| siRNA against p14 ARF Hs_CDKN2A_15 (SI02664403) | 5’-TACCGTAAATGTCCATTTATA-3’ | Qiagen,Venlo,Netherlands |
| siRNA against HIF-1α | 5’-AGGAAGAACTATGAACATAAA-3’ | Qiagen,Venlo,Netherlands |
| siRNA against HIF-2α | 5’-CCCGGATAGACTTATTGCCAA -3’ | Qiagen,Venlo,Netherlands |

**Supplementary Table S4:** List of the antibodies used in immunoblotting and immunofluorescence experiments.

| Antibody/Cat.number | Working dilution | Reference/Source |
|---|---|---|
| rabbit polyclonal anti-EXOSC10 (ab50558) | WB: 1/2500, IF: 1/500 | Abcam |
| mouse monoclonal anti- EXOSC10 (B-8: sc-374595) | WB: 1/1000, IF: 1/250 | Santa Cruz Biotechnology (Texas,USA) |
| mouse monoclonal anti-UBF (F-9:SC-13125) | WB: 1/1000, IF: 1/500 | Santa Cruz Biotechnology (Texas,USA) |
| rabbit polyclonal anti-DIS3 (14689-1-AP) | WB: 1/2000 | Proteintech |
| rabbit polyclonal anti- SKIV2L2 (12719-2-AP) | WB: 1/5000 | Proteintech |
| rabbit polyclonal anti-MPHOSPH6 (10695-1-AP) | WB: 1/2000 | Proteintech |
| rabbit polyclonal anti-C1D (10711-1-AP) | WB: 1/1000 | Proteintech |
| rabbit polyclonal anti-EXOSC3 (15062-1-AP) | WB: 1/2000 | Proteintech |
| rabbit polyclonal anti-USP36 (14783-1-AP) | WB: 1/1000, IF: 1/250 | Proteintech |
| mouse monoclonal anti-USP36 (68165-1-Ig) | WB: 1/1000 | Proteintech |
| mouse monoclonal anti-EXOSC2 (66099-1-Ig) | WB: 1/2000 | Proteintech |
| rabbit polyclonal anti-Phospho-p70 S6 Kinase (Thr389) (108D2) | WB: 1/1000 | Cell Signaling (Massachusetts, USA) |
| rabbit polyclonal anti-p14 ARF (E3X6D) | WB: 1/500 | Cell Signaling (Massachusetts, USA) |
| rabbit polyclonal anti- Fibrillarin (C13C3) | WB: 1/1000, IF:1/250 | Cell Signaling (Massachusetts, USA) |
| rabbit polyclonal anti- SENP3 (D20A10) | WB: 1/3000, IF: 1/500 | Cell Signaling (Massachusetts, USA) |
| rabbit polyclonal anti-HA-tag (C29F4) | WB: 1/5000 | Cell Signaling (Massachusetts, USA) |
| mouse monoclonal anti-β-Tubulin (D3U1W) | WB: 1/10000 | Cell Signaling (Massachusetts, USA) |
| mouse monoclonal anti-NPM1 (MAB937) | WB: 1/1000, IF:1/500 | Merck Millipore (Massachusetts, USA) |

| | | |
|---|---|---|
| affinity purified rabbit polyclonal anti- HIF-1α | WB 1/500, IF:1/200 | [5] |
| mouse monoclonal anti-HIF-1α (54/HIF1a 610959) | WB: 1/5000, IF:1/100 | BD Transduction Laboratories (New Jersey, USA) |
| rabbit polyclonal anti-HIF-2α (NB100-122) | WB: 1/2000 | Novus Europe |
| mouse monoclonal M2 clone anti-Flag-tag (F1804, 1:2000 dilution) | WB: 1/2500, IF:1/500 | Sigma-Aldrich (St Louis, MO) |
| affinity purified polyclonal anti-SUMO-1 | WB: 1/1000 | Kindly provided by Melchior Lab (ZMBH, Germany) |
| mouse monoclonal anti-SENP1 (C-12) | WB: 1/1000 | Santa Cruz Biotechnology (Texas,USA) |
| Mouse monoclonal anti-Ubiquitin (F-11) | WB: 1/1000 | Santa Cruz Biotechnology (Texas,USA) |
| Mouse monoclonal phosphoserine anibody (Clone 19/pSer, 612546) | WB: 1/500 | BD Transduction Laboratories (New Jersey, USA) |

**Supplementary Table S5:** Sequences of primers used in qPCR for the amplification of the indicated genes.

| oligonucleotides | Sequence | Source |
|---|---|---|
| CA9 For | 5'-TGGCTGCTGGTGACATCCTA-3' | Eurofins |
| CA9 Rev | 5'-TTGGTTCCCCTTCTGTGCTG-3' | Eurofins |
| SENP2 For | 5'- CAGTTCTCAAAGAAGTCAGATG-3' | Sigma-Aldrich (St Louis, MO) |
| SENP2 Rev | 5'-GTTTTGACTCCGTGATTTTG-3' | Sigma-Aldrich (St Louis, MO) |
| FOLR1 For | 5'-GAGGTTCTATGCTGCAGCCA-3' | Eurofins |
| FOLR1 Rev | 5'-CATTAGGGCCAGGCTAAGCA -3' | Eurofins |
| RPLP1 For | 5'- AAGCAGCCGGTGTAAATGTTGAGC -3' | Eurofins |
| RPLP1 Rev | 5'-CATTGCAGATGAGGCTCCCAATGT-3' | Eurofins |

### Supplementary Figures

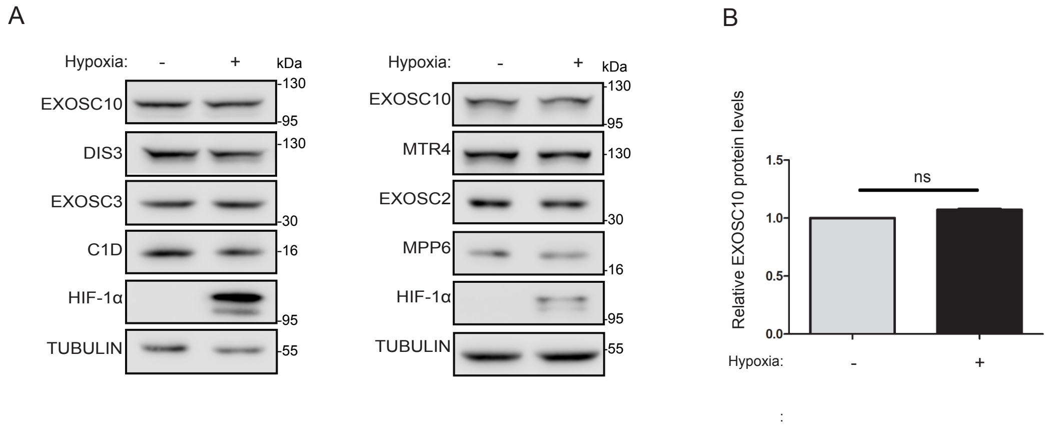


**Supplementary Figure S1: Hypoxia does not affect the expression levels of EXOSC10 or other RNA exosome components/cofactors. A.** HeLa cells were incubated in normoxia (-) or hypoxia (+) for 24 h and the endogenous expression levels of various RNA exosome components and cofactors were analyzed by immunoblotting using the indicated antibodies. Beta-tubulin was used as a loading control. **B.** Quantification of endogenous EXOSC10 expression levels (normalized to tubulin signal). Values are the mean of three independent experiments and are shown as fold increase compared to normoxic control (grey) and as mean ± standard error. For comparisons between groups unpaired t-test was used (n.s.: not significant).

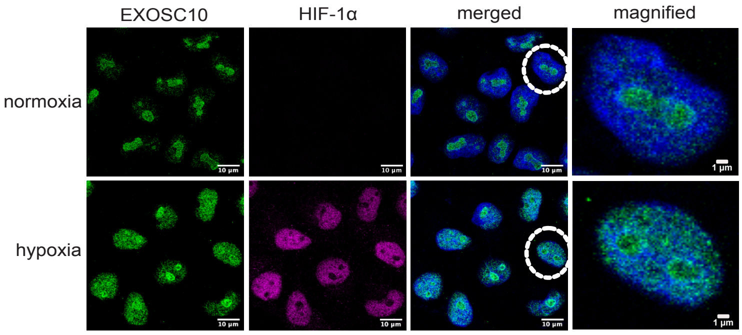


**Supplementary Figure S2: Hypoxia-dependent redistribution of EXOSC10 from the nucleolus to the nucleoplasm.** Indirect immunofluorescence analysis of HeLa cells incubated in normoxia or hypoxia using antibodies against EXOSC10 and HIF-1α (a marker for hypoxic conditions). DAPI was used for visualization of nuclear material. A cell, marked with dashed circle, is represented in magnified view as the last panel on the right for each case. Scale bar represents 10 µm or 1 µm as indicated.

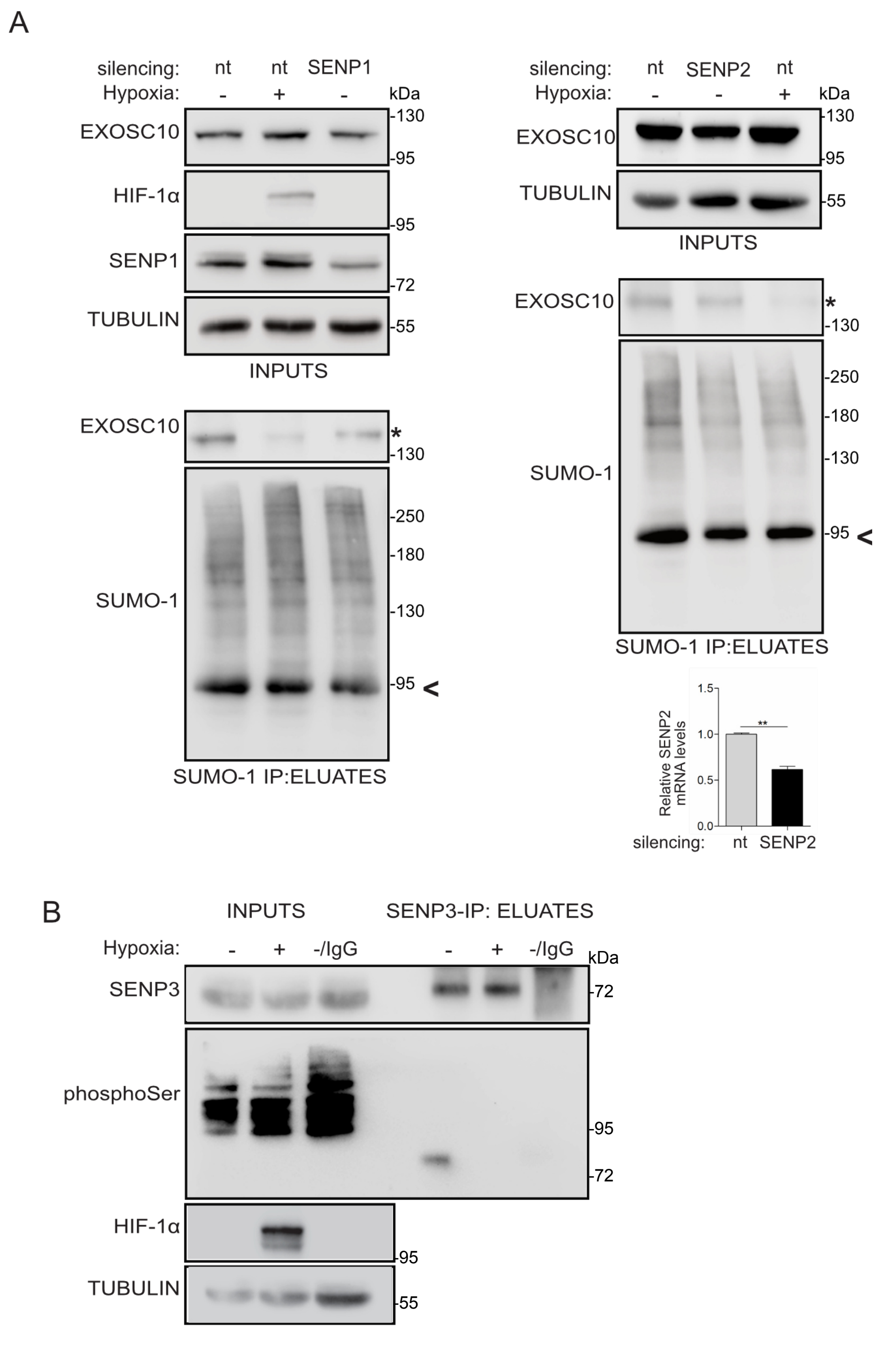


**Supplementary Figure S3: A. Silencing of SENP1 or SENP2 does not promote SUMOylation of EXOSC10.** HeLa cells were transfected with non-targeted siRNA (nt) or siRNAs against SENP1 (left) or SENP2 (right) and 24 h post-transfection, were incubated in normoxia (-) or hypoxia (+) for 24 h. Cell lysates were subjected to SUMO-1 IP. Inputs and eluates were analyzed by immunoblotting using the indicated antibodies. Silencing of SENP2 was verified with Q-PCR using specific primers for SENP2 mRNA. In all cases, the SUMOylated version of EXOSC10 is indicated with an asterisk (*). SUMOylated RanGAP1 is shown with an arrow and was indicated as a marker for equal loading in INPUTS (A) or as a marker for equal precipitation by anti-SUMO1. **B. SENP3 modulates its phosphorylation status in hypoxia.** HeLa cells incubated in normoxia (-) or hypoxia (+) for 24 h. Cell lysates were subjected to SENP3-IP. An equivalent amount of lysate was also incubated with IgG-beads and served as negative IP control. Soluble extracts (INPUTS) or anti-SENP3 and IgG immunoprecipitates (ELUATE) were analyzed by immunoblotting using the indicated antibodies. Beta-tubulin was used as loading control.

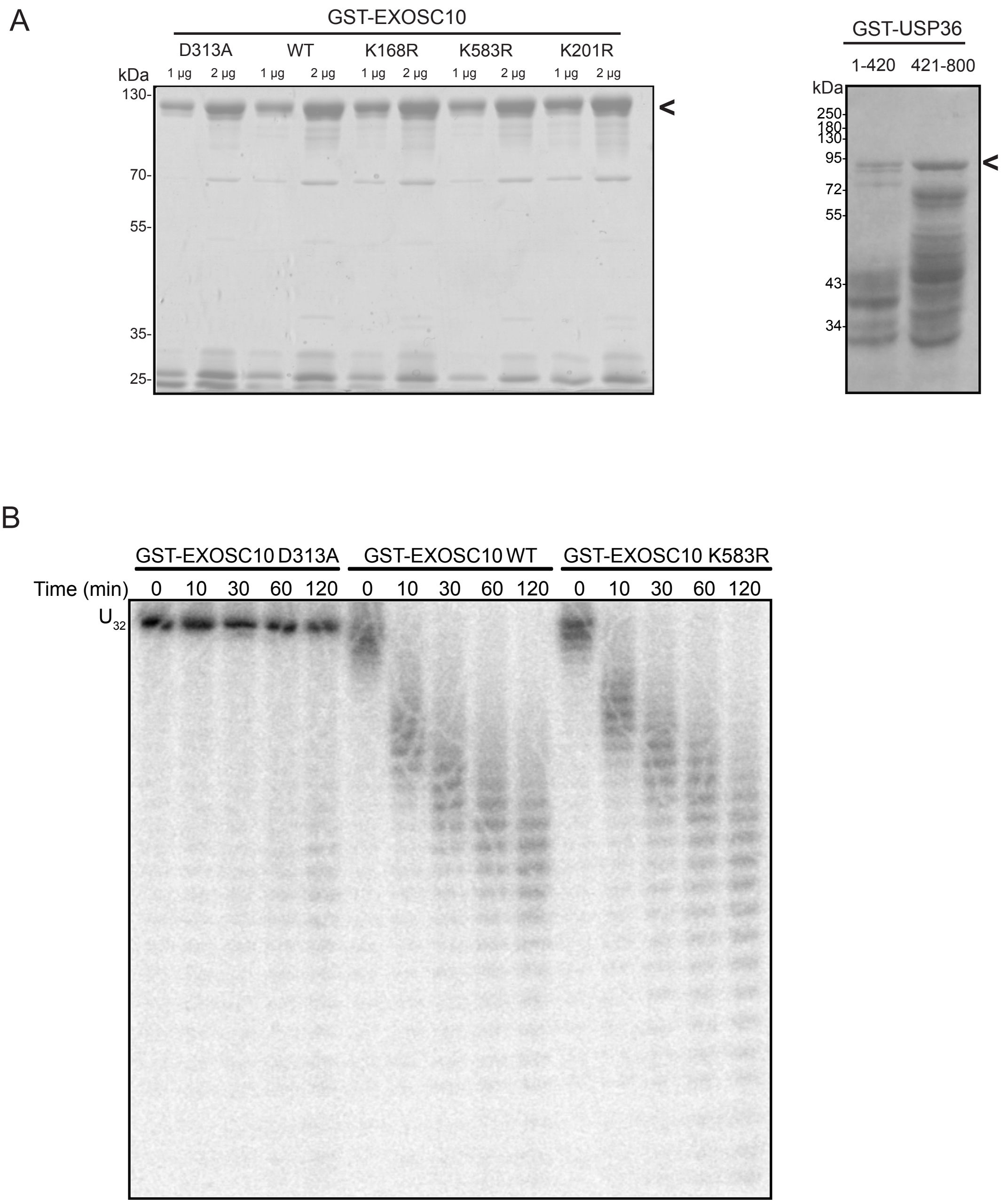


**Supplementary Figure S4: Purification and in vitro exoribonuclease assays of recombinant EXOSC10 forms. A.** Purified GST-tagged EXOSC10 WT and mutants D313A/K583R/K168R/K201R and GST-USP36 proteins (1-420 and 421-800 forms) were separated by SDS-PAGE and visualized by Coomassie staining. Position of MW is shown on the left. Full length purified proteins are shown with an arrowhead. **B.** *In vitro* exoribonuclease assays were performed using GST-tagged EXOSC10 WT, K583R or catalytically inactive D313A and [$^{32}$P]-labelled RNA (U32). Samples taken after the indicated times were separated on a denaturing (8 M urea) 12 % polyacrylamide gel and exposed to a phosphorimager screen before labelled RNAs and RNA fragments were visualized using a phosphorimager.

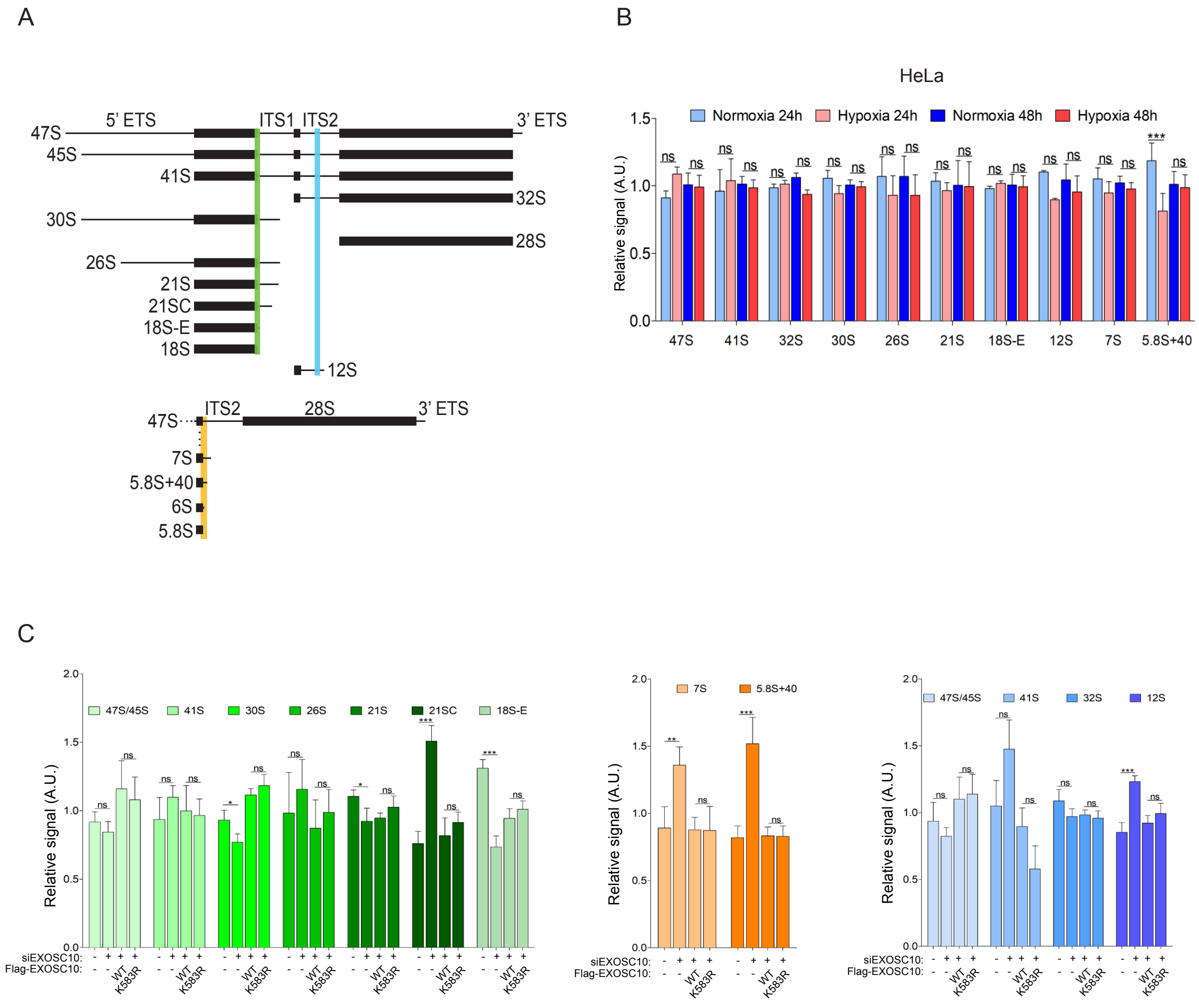


**Supplementary Figure S5: Pre-rRNA processing is not affected by deSUMOylation of EXOSC10. A.** The major pre-rRNA intermediates present in human cells are depicted schematically. Rectangles depict the mature 18S, 5.8S and 28S rRNAs and lines represent the internal transcribed spacer (ITS1 and ITS2) and the external transcribe spacer (5' ETS and 3' ETS). Colored lines indicate the base-pairing sites of probes used for northern blotting and the pre-rRNA intermediates detected. **B.** Quantification of the northern blot signals from Fig. 6C corresponding to the pre-rRNA intermediates in HeLa exposed to normoxia and hypoxia for 24 h and 48 h. The levels of pre-rRNA intermediates were obtained from three independent experiments, and the signals were quantified relative to the loading controls and shown as mean ± standard deviation. **C.** Total RNA extracted from stably transfected HeLa FlpIn cell lines expressing either the FLAG tag (-), or FLAG-EXOSC10 WT or FLAG-EXOSC10 K583R, in the presence of non-targeted siRNA (-) or a pool of siRNAs against EXOSC10 were analyzed by northern blotting as indicated in Fig. 6D. The levels of pre-rRNA intermediates detected with the probes 3'18S – 5'ITS1 (green), ITS2 (blue), and 3'5.8S – 5'ITS2 (orange) in three independent experiments were quantified relative to the loading controls and are shown as mean ± standard deviation. For high-molecular-weight pre-rRNAs (green and blue), mitochondrial 16S rRNA was used as loading control. For low-molecular-weight pre-rRNAs (orange), mitochondrial $tRNA^{Phe}$ was used as loading control.
